# Building Blocks of Understanding: Constructing a Reverse Genetics Platform for studying determinants of SARS-CoV-2 replication

**DOI:** 10.1101/2024.02.05.578560

**Authors:** Marco Olguin-Nava, Patrick Bohn, Thomas Hennig, Charlene Börtlein, Anne-Sophie Gribling-Burrer, Nora Schmidt, Neva Caliskan, Lars Dölken, Mathias Munschauer, Redmond P. Smyth

**Affiliations:** Helmholtz Institute for RNA-based Infection Research, Helmholtz Centre for Infection Research, 97080 Würzburg, Germany; Institute of Virology and Immunology, University of Würzburg, 97080 Würzburg, Germany; Faculty of Medicine, University of Würzburg, 97080 Würzburg, Germany; Institute of Medical Virology, Goethe-University Frankfurt, 60596 Frankfurt am Main, Germany

## Abstract

To better understand viral pathogenesis, host-virus interactions, and potential therapeutic interventions, the development of robust reverse genetics systems for SARS-CoV-2 is crucial. Here, we present a reverse genetics platform that enables the efficient manipulation, assembly, and rescue of recombinant SARS-CoV-2. The versatility of our reverse genetics system was demonstrated by generating recombinant SARS-CoV-2 viruses. We used this system to generate N501Y and Y453F spike protein mutants. Characterization studies revealed distinct phenotypic effects, impact on viral fitness, cell binding, and replication kinetics. We also investigated a recently discovered priming site for NSP9, which is postulated to produce a short RNA antisense leader sequence. By introducing the U76G mutation into the 5’UTR, we show that this priming site is necessary for the correct production of genomic and subgenomic RNAs, and also for efficient viral replication. In conclusion, our developed reverse genetics system provides a robust and adaptable platform for the efficient generation of recombinant SARS-CoV-2 viruses for their comprehensive characterization.

**Significance statement:** In this study, we present a versatile reverse genetics platform facilitating the efficient manipulation, assembly, and rescue of recombinant SARS-CoV-2. Demonstrating its adaptability, we successfully engineered N501Y and Y453F spike protein mutants, each exhibiting distinct phenotypic effects on viral fitness, cell binding, and replication kinetics. We also investigated a novel negative sense priming site for NSP9, demonstrating a role in RNA production and viral replication. This straightforward reverse genetic system is therefore a powerful tool to generate recombinant viruses for advancing our understanding of SARS-CoV-2 biology.

## Introduction

Severe acute respiratory syndrome coronavirus 2 (SARS-CoV-2) is the causative agent of the respiratory illness COVID-19, which emerged in December 2019 and has since unleashed a pandemic that continues to impact human health^1^. The virus contains a large (30 kb), capped and polyadenylated positive sense genome comprised of numerous open reading frames (ORFs) and flanked by two terminal untranslated regions (UTRs). The 5’ and 3’ UTRs are short but highly structured^2,3^. The 5’-terminal region contains two overlapping ORFs, ORF1a and ORF1b, which encode for two polyproteins, pp1a and pp1ab, that are cleaved into 16 non-structural proteins (NSPs) with multiple activities required for viral infection^3,4^. Pp1ab is translated using a minus 1 programmed ribosomal frameshift event to produce the RNA-dependent RNA polymerase (RdRp). Meanwhile, the 3′-terminal region (one-third of the genome) encodes the structural proteins which are expressed from subgenomic RNAs^5^. Like many RNA viruses, SARS-CoV-2 exhibits rapid rates of evolution^6–9^. Mutations arising in the SARS-CoV-2 genome, especially in “variants of concern”, have been found to impact infectivity, transmissibility or evasion of the immune system compared to the original strain^7,10,11^. It is because of this, and even with the availability of highly effective vaccines^12–14^, that the epidemic continues to spread rapidly around the world.

Molecular virology is a powerful approach for the characterization of viral evolution, pathogenesis, replication and immunogenicity. For RNA viruses, so called ‘reverse genetics’ (RG) enable the generation of recombinant viruses to investigate their pathogenic properties, including their interactions with the host cell^15–17^. Reverse genetics systems can also be used to study the behavior of mutant viruses in antiviral drug screening or therapeutic strategies^18–20^. They can also be used in diagnostics^21–24^, or to generate attenuated viruses for use as vaccines^15,25–27^. The first RG system for an RNA virus was developed for bacteriophage Qbeta by cloning full-length cDNA into one plasmid^28^. Following this plasmid transformation into E. coli, plaques were formed through infectious viruses. Subsequent work showed that this general approach could be applied to a variety of DNA and RNA viruses^29–33^.

Unfortunately, RG systems for viruses with large genomes, such as that of SARS-CoV-2, have proven technically challenging to develop, mainly because plasmids containing large viral inserts are unstable or toxic in *E. coli*^16^. Bacterial Artificial Chromosome (BAC)^23,34^ or Yeast Artificial Chromosome (YAC)^35,36^ are high-capacity vectors that allow the propagation of very large sequences in bacteria and yeast, respectively. Both YAC and BAC vectors have been purposed for the rescue of SARS-CoV-2 infectious molecular clones^18,19,23,37^. Although such systems allow for the generation of recombinant SARS-CoV-2 viruses, they present some disadvantages in that they could be challenging to construct, and subsequent genetic manipulation is more arduous than equivalent small high or low copy plasmids. Alternatively, SARS-CoV-2 can be split into smaller DNA fragments for later assembly either *in vitro* or in cells^22,38–43^. Genome splitting has the advantage that plasmids can be genetically manipulated using widely established techniques, but DNA fragments must be later assembled with high fidelity to form a contiguous genome. Some of these systems require RNA production to be transfected or electroporated into cells, involving an extra step that could lead to challenges or difficulties. Scarless and directional assembly has been achieved with Type IIS restriction enzymes that cleave outside of their recognition sequences, so called golden gate assebmly^22,38–40^. Another strategy, which is PCR based, is circular polymerase extension reaction (CPER) ^41,42,44^. It has even been reported that assembly may take place spontaneously in cells, provided the individual fragments have extensive overlaps^43^.

Here, we established a SARS-CoV-2 reverse genetics system based on 6 individual fragments assembled by PCR (**Figure 1**). We cloned the genome into 6 plasmids allowing its stable propagation and its genetic manipulation with well-established cloning methodologies^45^. PCR assembled fragments are then transfected into mammalian cells using polyethylenimine (PEI), a low-cost and easily accessible reagent. We used this strategy to generate two recombinant SARS-CoV-2 spike mutant viruses, Y453F and N501Y, which were first identified in variants of concern. In addition, we generated a recombinant virus containing the non-coding U76G mutation in the 5’UTR of the SARS-CoV-2 genome. The recombinant viruses were characterized, showing distinct phenotypic effects when tested for replication, cell binding, and subgenomic mRNA synthesis.

**Figure 1.**
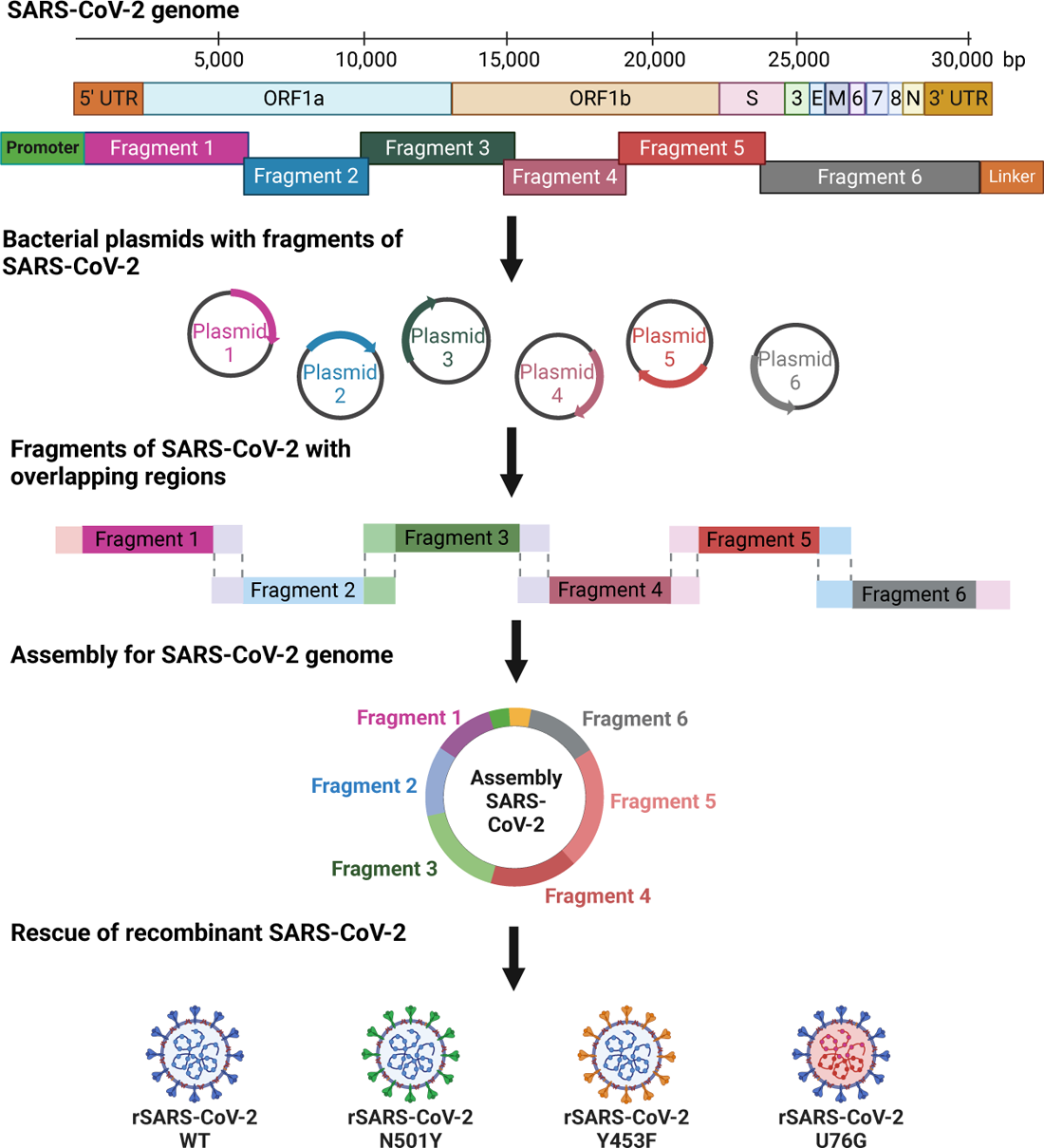
General overview of Reverse genetics system for SARS-CoV-2. SARS-CoV-2 is split in six fragments and cloned into bacterial plasmids. After PCR reaction, each fragment has 20 bp overlapping ends that allow the assembly of the complete viral genome. Then, SARS-CoV-2 assembly is transfected into mammalian cells in order to rescue recombinant virus.

Overall, the presented reverse genetics system can be used to explore the functional effects of nucleotide level mutations on SARS-CoV-2 replication.

## Results

### Dividing a long viral genome

The 30 kb genome of SARS-CoV-2 is one of the largest of any known RNA virus, making its handling and genetic manipulation an important technical challenge^3,17,18^. Therefore, we decided to divide the genome into smaller regions that can be individually propagated in *E. coli*. In this way, the maintenance of plasmids does not require special bacterial strains or protocols, each fragment can be modified more easily, and a variety of recombinant mutant viruses can be assembled by mixing different plasmids containing the desired mutations across the genome.

Our first approach consisted of splitting the full SARS-CoV-2 genome into six fragments to clone them in a bacterial vector. Fragments from 6.2 to 3.2 kb were amplified using as template the Yeast Artificial Chromosome (YAC) pCC1BAC-HIS3-SARS-CoV-2 containing the sequence of SARS-CoV-2 isolate Wuhan-Hu-1 (GenBank MN996528.3). After the generation of six fragments, one of them (3.5 kb) was cloned into the high copy number plasmid (Puc19) meanwhile the reminding five (6.2 to 3.2 kb) hard-to-clone fragments were cloned into pUA66, which is a low copy number plasmid to improve stability. An extra fragment (F6 GFP) which contains a reporter Turbo GFP sequence instead of the ORF7 located in the 3’ region of the genome was amplified using the YAC pCC1BAC-HIS3-SARS-CoV-2 GFP (BioProject: PRJNA615319; BioSample: SAMN14450690; Sample name: GFP-2_rSARS-CoV-2) as template and cloned into pUA66. All fragments were then subjected to Sanger sequencing. Except for a silent point mutation at position C28103A located in Fragment 6 WT, our sequence was identical to the original sequence of SARS-CoV-2 isolate Wuhan-Hu-1.

### Rebuilding a split SARS-CoV-2 genome

The next step in the establishment of the reverse genetics system was the reassembly of the full genome. As a first approach, we used a reassembly strategy based on Type IIs restriction enzymes (**Supplementary Figure 1A**). Type IIs enzymes recognize asymmetric DNA sequence outside the cleavage site producing specific and scarless ends that lead to directional cloning. This cloning strategy was used to reassemble the full viral genome involving a series of digestion/ligation cycles, similar to Golden Gate cloning^38,46^. Each fragment was amplified using a High-Fidelity DNA Polymerase with specific primers containing *Bsa*I sites. Subsequently, all fragments were individually digested using *Bsa*I for three hours at 37°C, followed by column purification (**Supplementary Figure 1B**). We decided to use *Bsa*I since two cleavage sites are already present in the genome. Therefore, extra *Bsa*I cleavage sites were engineered and designed for cloning and assembly purposes. Next, the fragments were ligated. For this, we initially mixed equimolar amounts of each digested fragment and incubated at 16°C and 4°C overnight (OVN) in the presence of T4 DNA ligase. The results showed the presence of multiple bands but not the expected 30 kb viral genome, demonstrating a partial and inefficient assembly (data not shown). Later, two different ligation strategies were tested to assemble the viral genome: 1) two reactions including fragments F1, F2 and F3 in the first reaction and fragments F4, F5 and F6 in a second reaction, and 2) three reactions containing F1 plus F2, F3 plus F4, and F5 plus F6. For both reactions, the ligation condition tested was 4°C OVN. Immediately after, the two or three reactions were mixed and incubated for an extra ligation step to assemble the full viral genome at 4°C OVN (**Supplementary Figure 1C**). Two additional temperatures, 16°C or room temperature (RT) OVN, for the final assembly were also tested (**Supplementary Figure 1D** and 1E). All the conditions tested lead to the desired ligation product of 30 kb, but the assembly was incomplete, with several intermediate products and an excess of residual unused fragments observed.

In light of the above, we decided to change the assembly strategy for a scheme based on circular polymerase extension reaction (CPER), which has been described before for SARS-CoV-2 and other RNA viruses^24,41,42,44,47^. This approach is based on the generation of the different fragments F1-F6 for rSARS-CoV-2 WT, all of them containing an overlap of 20 bp between adjacent fragments. Three plasmids were modified to adapt them to this method. First, a CMV promoter (598 bp) was added to the upstream region of the 5’UTR in fragment F1. This promoter allows the transcription of the viral genome by cellular polymerase II in the nucleus. Next, a linker sequence was introduced after the 3’UTR in fragments F6 WT and F6 GFP. The linker consists of three different elements: 1) a hepatitis delta virus ribozyme (HDVr) to produce a specific 3’ end in the viral RNA, 2) a simian virus 40 PolyA signal (SV40 polyA) to enhance transcription termination and to promote RNA stability, and 3) a spacer sequence of 364 bp to create an intermediate area between the extremes of the viral genome during the circularization assembly^44^ (**Supplementary Figure 2A**).

Following amplification and purification of the individual fragments (**Figure 2A**), each of them were mixed in an equimolar amount 0.1 pMol. During temperature cycling in the presence of a high-fidelity DNA polymerase and dNTPs, complementary fragments can align to create a circular full genome assembly. For the assembly of rSARS-Cov-2 WT or GFP, fragments F1-F5 and either the F6 WT or F6 GFP were used, respectively. The latter contains Turbo-GFP instead of ORF7. Initially, two cycling conditions were tested: 1) initial denaturation at 98°C for 30 sec, 12 cycles of 10 sec at 98°C, 20 sec at 55°C and 10 min at 68°C, and a final elongation for 12 min at 68°C. 2) initial denaturation at 98°C for 2 min, 20 cycles of 10 sec at 98°C, 15 sec at 55°C and 25 min at 68°C, and a final elongation for 25 min at 68°C. Only the first condition produced the full genome assembly of 30 kb (**Figure 2B**). Since the main difference between the two assembly programs consisted in the number of cycles (12 and 20) and the best condition was the one with the fewer cycles (12 cycles), we decided to further optimize the assembly reaction by testing 6 and 9 cycles. Both reactions produced incomplete assemblies with multiple bands indicating intermediate products and unused initial fragments (**Supplementary Figure 2B** and 2C).

**Figure 2.**
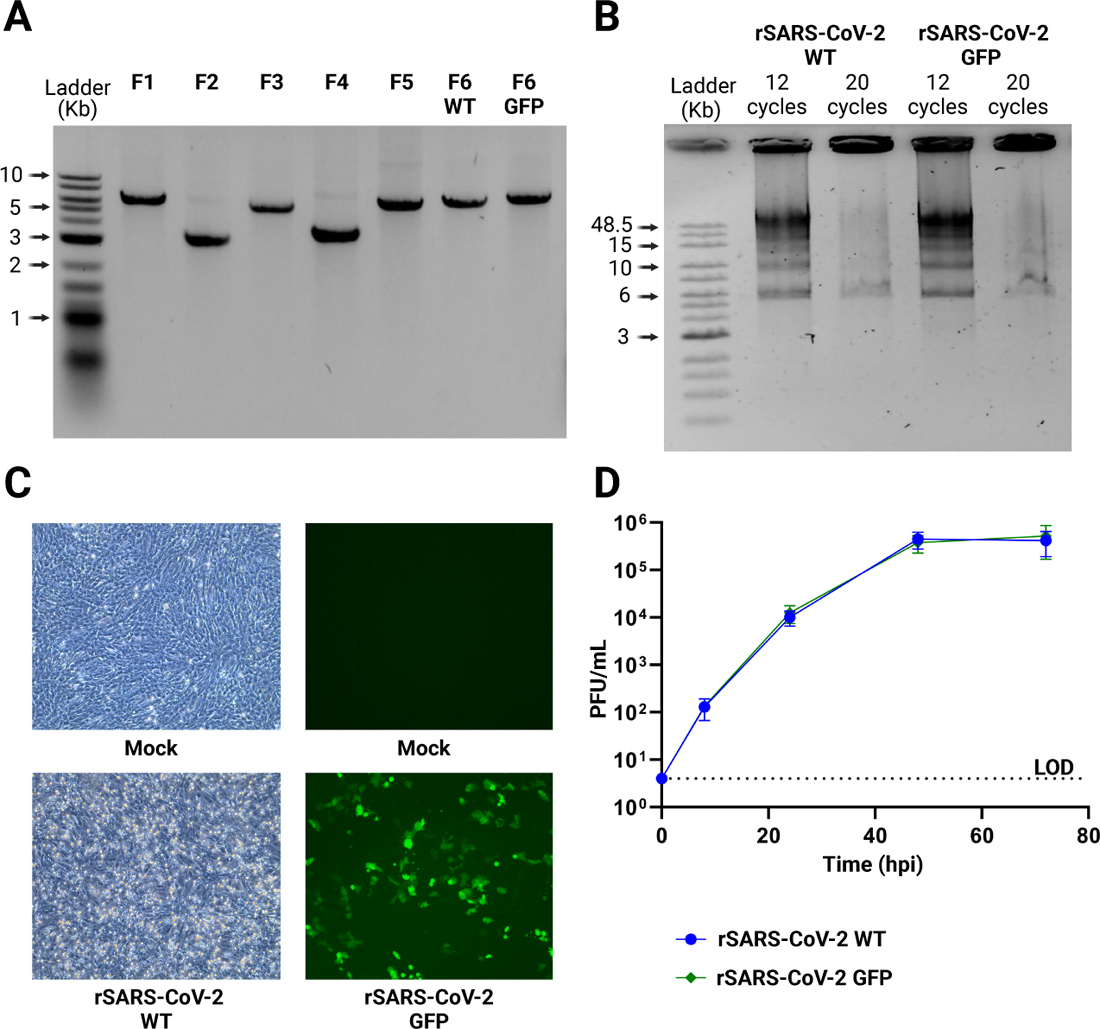
Rescue and characterization of recombinant SARS-CoV-2. **(A)** DNA fragments of SARS-CoV-2. PCR products of 7 fragments of SARS-CoV-2, including the fragment for the GFP reporter, using bacterial plasmids as template. **(B)** Assembly of complete DNA SARS-CoV-2. DNA fragments are assembled to produce the 30 kb SARS-CoV-2 DNA, two different conditions with different number of assembly cycles were tested. **(C)** Infection using recombinant SARS-CoV-2. After DNA transfection, supernatants were collected and used to infect a monolayer of Vero E6 TMPRSS2, after 2 days cells: present CPE for SARS-CoV-2 WT or express GFP for the recombinant SARS-CoV-2 GFP. **(D)** Growing kinetics of rSARS-CoV-2. Comparison of growth kinetics of rSARS-CoV-2 WT and rSARS-CoV-2 GFP in Vero E6 TMPRSS2 cells infected at a MOI 0.01 over a 3-day time course. LOD: Limit of detection. *n* = 2 independent experiments with two replicates each. The graphic shows mean values ± SD. Stars indicate the degree of significance compared to the rSARS-CoV-2 WT control condition (*=p<0.05, **= p<0.01, ***= p<0.001, ****= p<0.0001 by two-sided unpaired Student’s *t*-test).

Considering all the above, we decided to use a PCR-based assembly protocol consisting of 12 cycles as the main strategy to assemble the full genome of rSARS-CoV-2 WT and rSARS-CoV-2 GFP.

### Rescue of recombinant SARS-CoV-2

For the rescue of recombinant SARS-CoV-2, we opted to transfect the assembled DNA directly into different easy-to-transfect cells, as our system incorporates a CMV promoter capable of transcription by cellular machinery. The transfected cells were then plated onto SARS-CoV-2-susceptible cells followed by two further passages to amplify any infectious virus. Since the expression of the SARS-CoV-2 nucleoprotein (N) has been reported to enhance the rescue of virus^18,38^, a plasmid facilitating the expression of the N protein was co-transfected with the assembled SARS-CoV-2 DNA. Briefly, the assembly was reverse transfected into HEK293T or HEK293T ACE2 cells using a variety of transfection reagents, such as Lipofectamine 2000, Lipofectamine 3000, TransIT 2020, TransIT LT1 and polyethylenimine (PEI). After 24 h, transfected cells were detached with trypsin and seeded on top of a monolayer of VERO E6 TMPRSS2 cells, which are highly susceptible to infection and thus enhance the isolation of SARS-CoV-2. At 10 days post-transfection (dpt) supernatants were collected. Surprisingly, the only condition that yielded infectious virus was when HEK293T ACE2 cells were transfected using PEI. Using this method, we obtained infectious rSARS-CoV-2 WT and GFP, which caused significant cytopathic effects (CPE) or GFP expression (**Figure 2C**) and replicated to titers of 2.9 x10^5^ PFU/mL and 2.2 x10^5^ PFU/mL, respectively. Further preliminary analysis showed that plaque morphology or size were similar for both viruses (**Supplementary Figure 3**).

Next, we investigated the replication kinetics of the recombinant viruses rSARS-CoV-2 WT and rSARS-CoV-2 GFP. VERO E6 TMPRSS2 cells were infected for 8, 24, 48 and 72 hours post infection (hpi) using both recombinant viruses at a MOI of 0.01. Growth kinetics of both recombinant viruses were virtually identical demonstrating that the rescue of SARS-CoV-2 was successful and that the substitution of ORF7 for reporter GFP did not affect the replication of SARS-CoV-2 in Vero E6 TMPRSS2 cells during the period evaluated (**Figure 2D**).

In conclusion, efficient rescue of infectious rSARS-CoV-2 was achieved using a simple and economical transfection protocol utilizing PEI to transfect unpurified, assembled DNA in combination with a plasmid coding for N.

### Analysis of the full genome of recombinant SARS-CoV-2 using nanopore sequencing

To evaluate the accuracy of the reverse genetics system and the stability of the genome we sequenced recombinant viruses after two passages using Oxford Nanopore Sequencing (**Supplementary Figure 4A**). RNA of supernatants belonging to passage 1 and passage 2 of rSARS-CoV-2 WT and rSARS-CoV-2 GFP in Vero E6 TMPRSS2 were isolated, reverse transcribed and amplified as DNA amplicons. Fourteen amplicons of 2.4 kb using two panels of primers based on the Artic sequencing protocol for SARS-CoV-2 were generated (**Supplementary Figure 4B**). Afterwards, each of fourteen amplicons were purified, pooled, multiplexed, barcoded and sequenced using an Oxford Nanopore MinION device.

All the recombinant viruses were compared to the sequence of SARS-CoV-2 isolate Wuhan-Hu-1 (GenBank MN996528.3) or YAC pCC1BAC-HIS3-SARS-CoV-2 GFP (BioProject: PRJNA615319; BioSample: SAMN14450690). When examining mutations with a frequency exceeding 50%, the initial passage of the four-rescued rSARS-CoV-2 WT demonstrated a nucleotide accuracy of 99.9933% (**Table 1**). Subsequently, we subjected one of the four-rescued rSARS-CoV-2 WT to a second passage in VERO E6 TMPRSS2, revealing a nucleotide accuracy of 99.9866% upon sequencing. Similarly, both the first and second passage of rSARS-CoV-2 GFP showed high fidelity rescue, with a nucleotide accuracy of 99.9967% (**Table 1**).

**Table 1.**
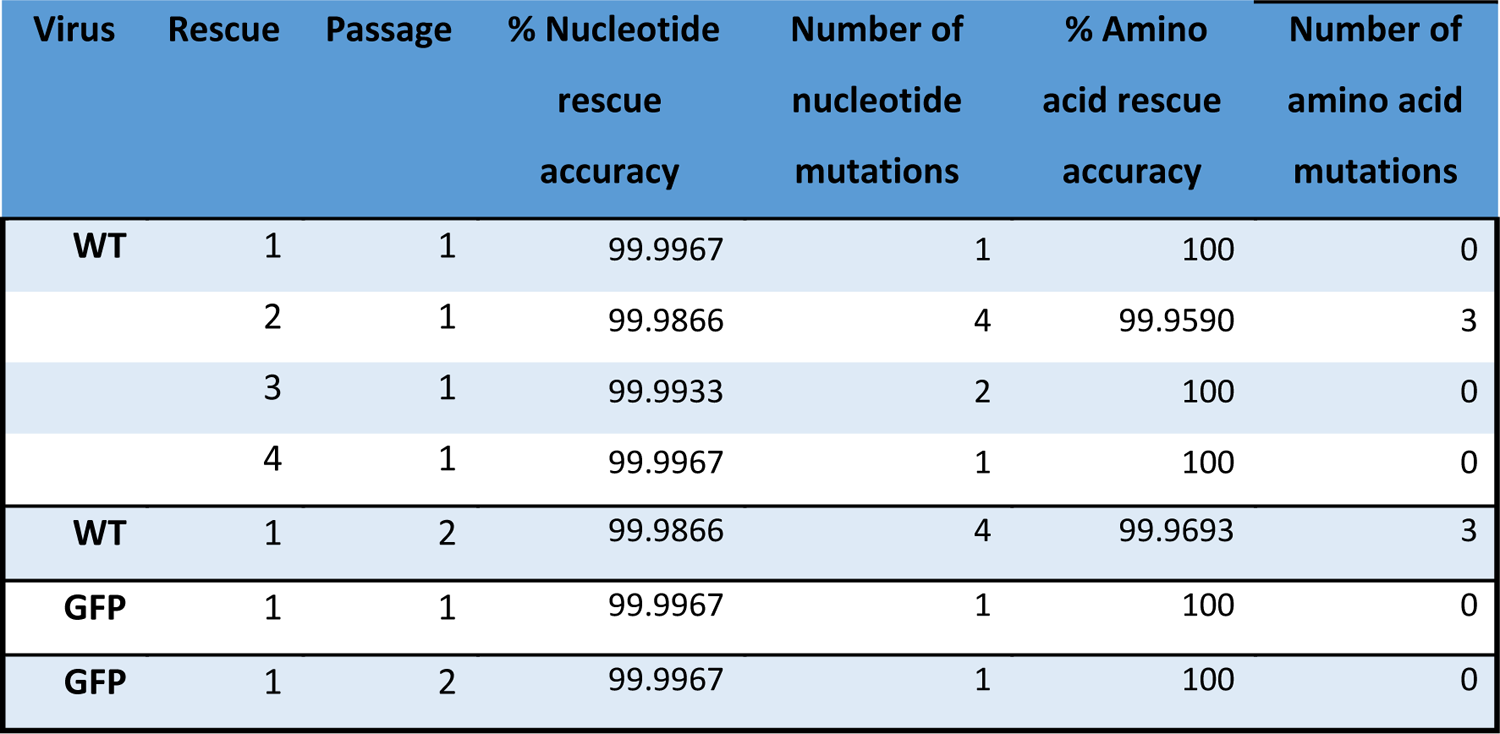
Rescue accuracy of SARS-CoV-2.

At the amino acid level, we identified four non-synonymous mutations (P23L and A1527V in NSP3, I382V in NSP4 and T71M in N) for one of the rSARS-CoV-2 WT isolates at the first passage, giving a fidelity of 99.959%. In a different rSARS-CoV-2 WT isolate, we did not identify any non-synonymous mutations in the first passage, but three arose in the second passage (T1180S and A1305G in NSP3 and A701T in S), suggesting potential selection for mutations during passaging, although still with an amino acid accuracy of 99.969% (**Figure 3**, **Table 1** and **Supplementary Table 1**). No mutations were detected in rSARS-CoV-2 GFP in both passages (**Figure 3**).

**Figure 3.**
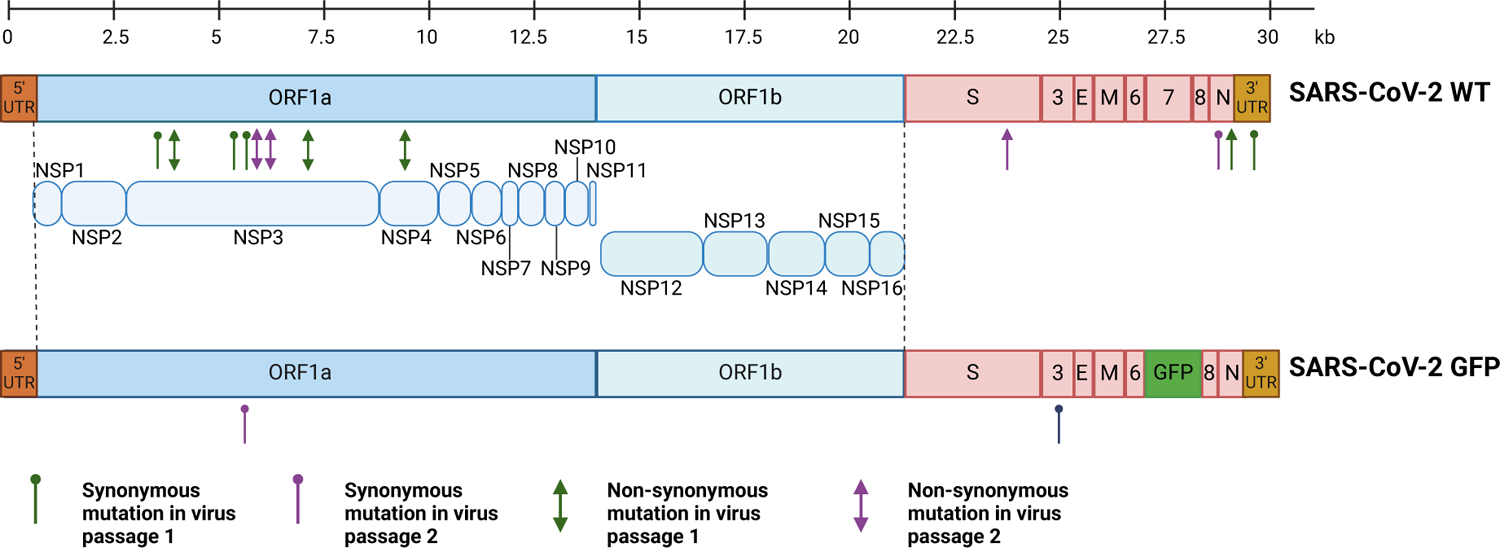
Sequencing of rSARS-CoV-2 using Oxford Nanopore sequencing. Schematic representation of rSARS-CoV-2 WT and rSARS-CoV-2 GFP genome. Synonymous *(hairpins)* and non-synonymous *(arrows)* mutations of each virus are indicated, specifying those mutations present in two different passages (*passage 1=green* and *passage 2=purple*).

Collectively, our reverse genetics system demonstrated to be able to rescue rSARS-CoV-2 with a consistent accuracy of over 99.9% for both nucleotide and amino acid sequence.

### Generation of SARS-CoV-2 spike mutants

The transmembrane spike glycoprotein is responsible for SARS-CoV-2 entry into the host cell. Initially, the spike receptor-binding domain (RBD) binds to the human angiotensin-converting enzyme 2 (ACE2) receptor to start the infection process (**Supplementary Figure 5A**). The spike protein is exposed to the humoral immune system on the surface of viral particles, and the encoding region contains the highest number of non-synonymous mutations in the entire SARS-CoV-2 genome^7,48,49^. Spike mutations are highly relevant because they are responsible for influencing the virus host range, tissue tropism, and pathogenesis. Most experiments testing the impact of this mutations have been performed only using purified proteins with *in vitro* studies or pseudovirus expressing the spike protein, thus lacking the presence of other factors or interactions between the cell and the virus that could influence the natural course of the infection^50–57^. We chose to evaluate two spike mutations, N501Y (Alpha, Beta, Gamma and Omicron variants) and Y453F (Cluster 5 variant), both associated with an increase in ACE2-binding affinity and located in the RBD-ACE2 interface^7,5^>.

Four recombinant viruses based on rSARS-CoV-2 WT or GFP, and containing Y453F or N501Y spike mutations were created to evaluate their importance in the viral cycle. First, plasmid F5 was modified by Direct Site Mutagenesis^45^ to introduce A22920T and A23053T exchanges for Y453F and N501Y, respectively (**Supplementary Figure 5B**). For the assembly, we followed the same conditions as before (**Supplementary Figure 5C** and 5D). We obtained preparations for each of the recombinant isolates with titers of 3.4 x10^5^ PFU/mL for rSARS-CoV-2 Y453F, 2) 5.0 x10^5^ PFU/mL for rSARS-CoV-2 N501Y, 3) 1.7 x10^6^ PFU/mL for rSARS-CoV-2 GFP Y453F and 4) 4.2 x10^5^ PFU/mL for rSARS-CoV-2 GFP N501Y. The successful introduction of each mutation was confirmed in the four recombinant viruses by RT-PCR followed by Sanger sequencing (**Figure 4A**). For all the recombinant viruses, CE of GFP expression was observed (**Figure 4B**) and no differences in shape or size of plaques were observed (**Supplementary Figure 3**).

**Figure 4.**
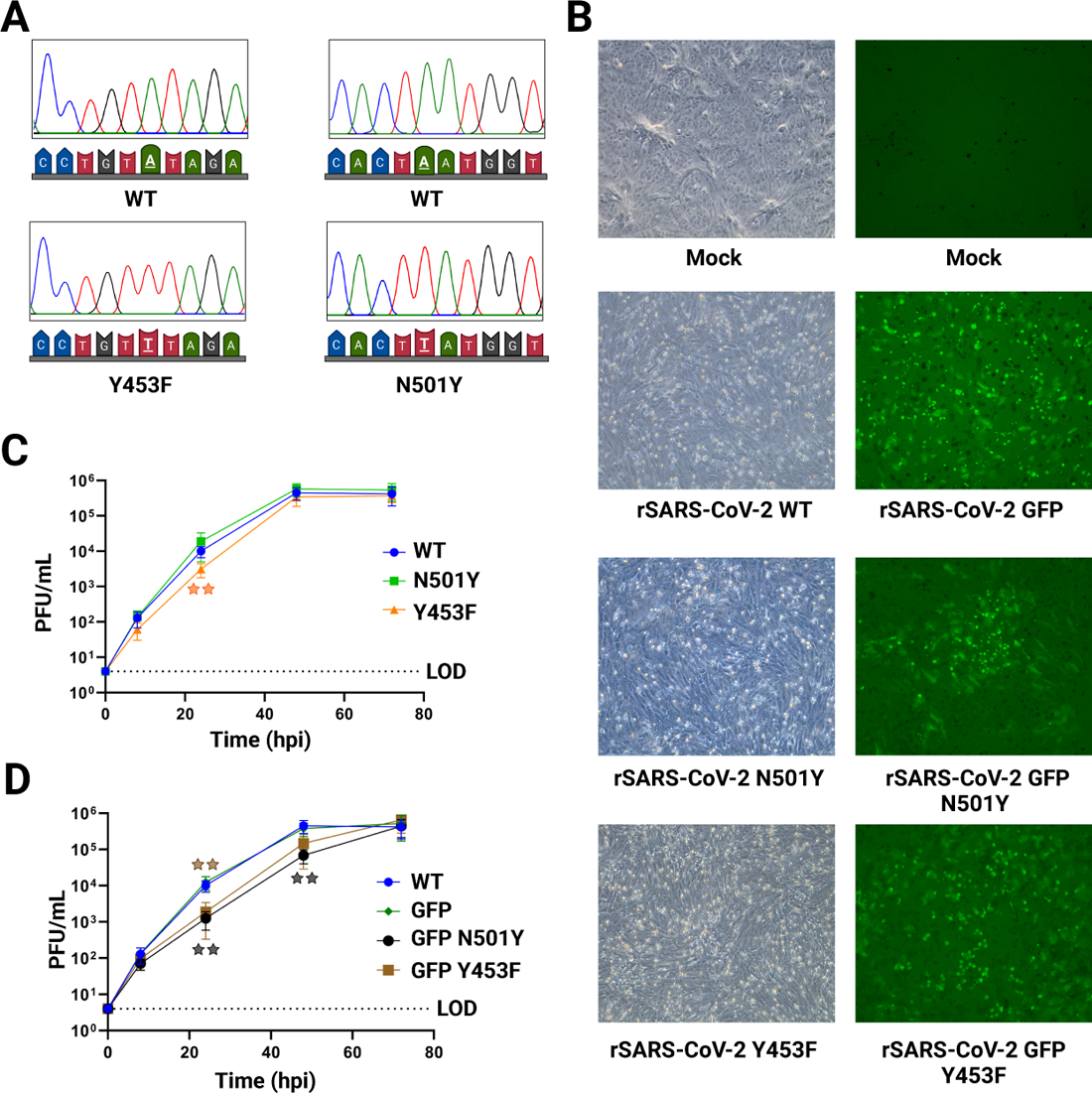
Characterization of SARS-CoV-2 spike mutants. **(A)** Sanger sequencing of SARS-CoV-2 RBD spike mutants. Supernatant of rSARS-CoV-2 N501Y, rSARS-CoV-2 Y453F, rSARS-CoV-2 GFP N501Y and rSARS-CoV-2 GFP Y453F infected cells were subjected to Sanger sequencing, showing the desired mutations compared to rSARS CoV-2 WT. **(B)** Infection using recombinant SARS-CoV-2 RBD spike mutants. After DNA transfection, supernatants were collected to infect a monolayer of Vero E6 TMPRSS2 cells, after 2 days CPE was showed for rSARS-CoV-2 WT, rSARS-CoV-2 Y453F and rSARS-CoV-2 N501Y; or reporter GFP was expressed in the case of rSARS-CoV-2 GFP, rSARS-CoV-2 GFP Y453F and rSARS-CoV-2 GFP N501Y. **(C)** Growth kinetics of rSARS-CoV-2 N501Y and Y453F. Comparison of growth kinetics of SARS-CoV-2 RBD spike mutant to rSARS-CoV-2 WT and rSARS-CoV-2 GFP in Vero E6 TMPRSS2 cells infected at a MOI 0.01 over a 3-day time course. **(D)** Growth kinetics of rSARS-CoV-2 GFP N501Y and Y453F. Comparison of growth kinetics of SARS-CoV-2 RBD spike mutant to rSARS-CoV-2 WT and rSARS-CoV-2 GFP in Vero E6 TMPRSS2 cells infected at a MOI 0.01 over a 3-day time course. *n* = 2 independent experiments with two replicates each. The graphic shows mean values ± SD. Stars indicate the degree of significance compared to the rSARS-CoV-2 WT or rSARS-CoV-2 GFP control (*=p<0.05, **= p<0.01, ***= p<0.001, ****= p<0.0001 by two-sided unpaired Student’s *t*-test).

Next, we characterized the mutant rSARS-CoV-2 viruses, evaluating their replication kinetics (**Figure 4C** and **4D**). Each virus was used to infect VERO E6 TMPRSS2 at a MOI of 0.01 and samples were collected at 8, 24, 48 and 72 hpi for titration by plaque assay. Compared to rSARS-CoV-2 WT, both spike mutants grew to virtually identical titers at 72 hpi with similar kinetics although the rSARS-CoV-2-Y453F showed a modestly reduced titer at 24 hpi. Although, the rSARS-CoV-2 GFP-derived spike mutants appeared to be delayed in replication, they reached similar peak titers compared to rSARS-CoV-2 GFP at 72 hpi.

Altogether, we successfully rescued four different viruses with mutant Spikes, proving our system is suitable for the introduction of point mutations in recombinant viruses. Both these mutations exhibit similar replication kinetics compared to rSARS-CoV-2 WT.

### Binding and competition assay for spike mutant SARS-CoV-2

We next explored the impact of the two RBD spike mutants on binding to the cell receptor ACE2. ACE2 is the main cell receptor for several coronaviruses including SARS-CoV, MERS-CoV and SARS-CoV-2^1,58,59^. For these studies, we used two different cell lines: Vero E6 cells overexpressing TMPRSS2 and A549 overexpressing ACE2. Vero E6 are susceptible to SARS-CoV-2 infection due to the presence of ACE2 receptor in their membrane but the overexpression of TMPRSS2 facilitates the virus entrance^59,60^. On the other hand, A549 cells are epithelial lung cells poorly infectable by SARS-CoV-2 due to the absence of the receptor ACE2^59^, but the overexpression of the ACE2 receptor leads to a susceptible cell line^61,62^. We speculated that the differential expression of ACE2 receptor in those cell lines could show a distinct phenotype during binding assays for the mutant viruses.

Binding assays are based on the incubation of a monolayer of cells with a virus at 4°C temperature, during which the interaction between the spike and the cell receptor can occur, but viral entry is inhibited as a result of the low temperature^63,64^. Vero E6 TMPRSS2 and A549 ACE2 cell lines were incubated with three recombinant SARS-CoV-2 viruses: rSARS-CoV-2 WT, rSARS-CoV-2 Y453F or rSARS-CoV-2 N501Y using MOI of 0.1 or 0.01 at 4°C for 1 h. Unbound virus particles were washed away with ice cold PBS and viral RNA from viruses attached to the cell monolayer were extracted. We compared the binding of each RBP spike mutant to ACE2 against rSARS-CoV-2 WT using RT-qPCR against the *RdRp* gene (**Supplementary Figure 6**). Interestingly, there was no difference in the binding efficiency between rSARS-CoV-2 WT and rSARS-CoV-2 N501Y in both cell lines using MOI of 0.1 (**Figure 5A** and **5B**). On the other hand, rSARS-CoV-2 Y453F showed a lower binding efficiency than rSARS-CoV-2 WT in both cell lines with an MOI of 0.1 (**Figure 5A** and **5B**). We found similar results when a MOI of 0.01 was used (**Supplementary Figure 6B** and 6C).

**Figure 5.**
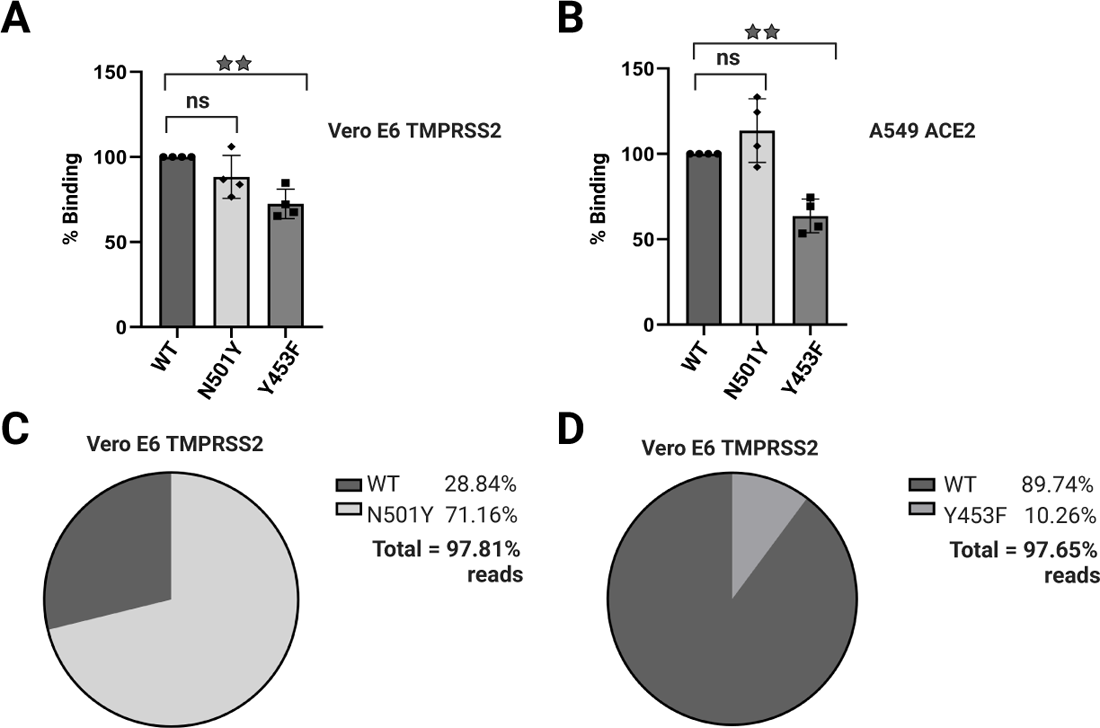
Binding and competition assays for recombinant SARS-CoV-2 RBD spike mutants. **(A** and **B)** Binding assay for SARS-CoV-2 spike mutants. Vero E6 TMPRSS2 cells or A549 ACE2 were incubated with rSARS-CoV-2, rSARS-CoV-2 Y453F or rSARS-CoV-2 N501Y at 4°C to evaluate the interaction between the spike and ACE2 receptor. Samples were evaluated by qPCR for *RdRp* and compared to rSARS-CoV-2 WT in **(A)** Vero E6 TMPRSS2 at a MOI 0.1 and **(B)** A549 ACE2 at a MOI 0.1. Quantification relative to 18S rRNA. *n* = 3 independent experiments with two replicates each. The graphic shows mean values ± SD. Stars above the bars indicate the degree of significance compared to the control condition (*=p<0.05, **= p<0.01, ***= p<0.001, ****= p<0.0001 by one-way ANOVA). **(C** and **D)** Competition assay for SARS-CoV-2 spike mutants. Vero E6 TMPRSS2 were infected with rSARS-CoV-2 WT plus rSARS-CoV-2 N501Y and rSARS-CoV-2 WT plus rSARS-CoV-2 Y453F at a MOI 0.1. After 48 hpi samples coming from cells infected with **(C)** WT+N501Y or **(D)** WT+Y453F were collected and sequenced by Oxford Nanopore. Percentage of reads corresponding to the respective rSARS-CoV-2 mutant sequence. *n* = 2 independent experiments with two duplicates.

We next evaluated whether the introduced RBD spike mutants could give an advantage or disadvantage in the infectivity of SARS-CoV-2. We therefore performed competition assays in which Vero E6 TMPRSS2 or A549 ACE2 were co-infected at a MOI 0.1 with rSARS-CoV-2 WT plus rSARS-CoV-2 Y453F, or rSARS-CoV-2 WT plus rSARS-CoV-2 N501Y. As controls, we also infected both cell lines with rSAR-CoV-2 WT, rSARS-CoV-2 Y453F and rSARS-CoV-2 N501Y alone. After 48 h, RNA from cells was extracted and prepared for sequencing on an Oxford Nanopore MinION flowcell, as described before (**Supplementary Figure 7A**). In cells infected with only one virus, we observed a correlation of over 90% between each recombinant virus and its corresponding sequence (**Supplementary Figure 7B** and C). The competition assays disclosed interesting phenotypes for rSARS-CoV-2 N501Y depending on the cell line evaluated. In Vero E6 TMPRSS2 cells, the N501Y mutant overtook WT with 71.16% presence versus 28.84% inside the cell (**Figure 5C**). In contrast, in the competition assay conducted on A549 ACE2, the distribution was similar between WT with 50.25% and 47.97% of N501Y (**Supplementary Figure 7D**). On the other hand, rSARS-CoV-2 WT outperformed the Y453F mutant in both cell lines with 89.79% against 10.26% in Vero E6 TMPRSS2, and 79.24% against 17.49% in A549 ACE2 respectively (**Figure 5D** and **Supplementary Figure 7E**).

Taken together, our results reveal distinct phenotypes for the N501Y and Y453F mutants in terms of ACE2 binding and viral infectivity. While N501Y exhibited comparable binding efficiency to WT in both cell lines, its performance in competition assays revealed cell-line-specific outcomes. For the Y453F mutant exhibited decreased binding compared to the WT virus, which likely contributed to its lower fitness in competition assays.

### A role for anti-leader sequence in SARS-CoV-2 replication

During the replication cycle of SARS-CoV-2, a set of subgenomic RNAs (sgRNA) are generated to produce viral structural and non-structural proteins. sgRNAs synthesis starts at the 3’ end of the genome and employs a discontinuous transcription mechanism to generate the negative-sense sgRNAs, which are later transcribed into positive-sense subgenomic mRNAs (sgmRNAs)^2,5^. Template-switching is proposed as the main mechanism for discontinuous transcription, which requires a 50 bp leader transcription regulatory sequence (TRS-L) in the 5’UTR and eight body transcription regulatory sequences (TRS-B) located across the genome. Although the molecular mechanism is incompletely understood, a base pairing interaction of the TRS-L with the TRS-B upstream of each gene presumably leads to fusion of the leader-body to generate at least eight sgRNAs, each of which possess the same initial 5’ and 3’ ends^2,3,5,65,66^.

Recently NPS9 has been reported to be involved in priming of RNA synthesis^67–72^. A recent study used covalent RNA immunoprecipitation (cRIP) to show that NSP9 is covalently attached to the 5’ ends of SARS-CoV-2 RNAs ^70^. cRIP is a variant of crosslinking immunoprecipitation (CLIP) without the UV crosslinking step. In addition to the annotated genome ends, cRIP identified an unusual covalent linkage at position 76 in the negative-sense viral RNA. Interestingly, this site is located adjacent to the T-LTR, which suggests that NSP9 may prime anti-leader synthesis at this position, putatively influencing discontinuous transcription or template switching. To test this, we introduced a T76G point mutation into the SARS-CoV-2 5’UTR^45^ (**Supplementary Figure 8A**). Two assemblies for rSARS-CoV-2 U76G and rSARS-CoV-2 GFP U76G were generated following the same conditions as before (**Supplementary Figure 8B** and 8C). The two recombinant viruses were rescued producing CPE or GFP expression after 48 hpi in Vero E6 TMPRSS2 cells (**Supplementary Figure 8D**). Titration of the viral stocks by plaque assay indicated similar rescue efficiencies of 7.0 x10^5^ PFU/mL for rSARS-CoV-2 U76G and 5.4 x10^5^ PFU/mL for rSARS-CoV-2 GFP U76G. Notably, the U76G mutant viruses had a reduced plaque size compared to rSARS-CoV-2 WT (**Supplementary Figure 3**). Each virus was analyzed using Sanger sequencing to demonstrate the presence of the intended mutation (**Figure 6A**).

**Figure 6.**
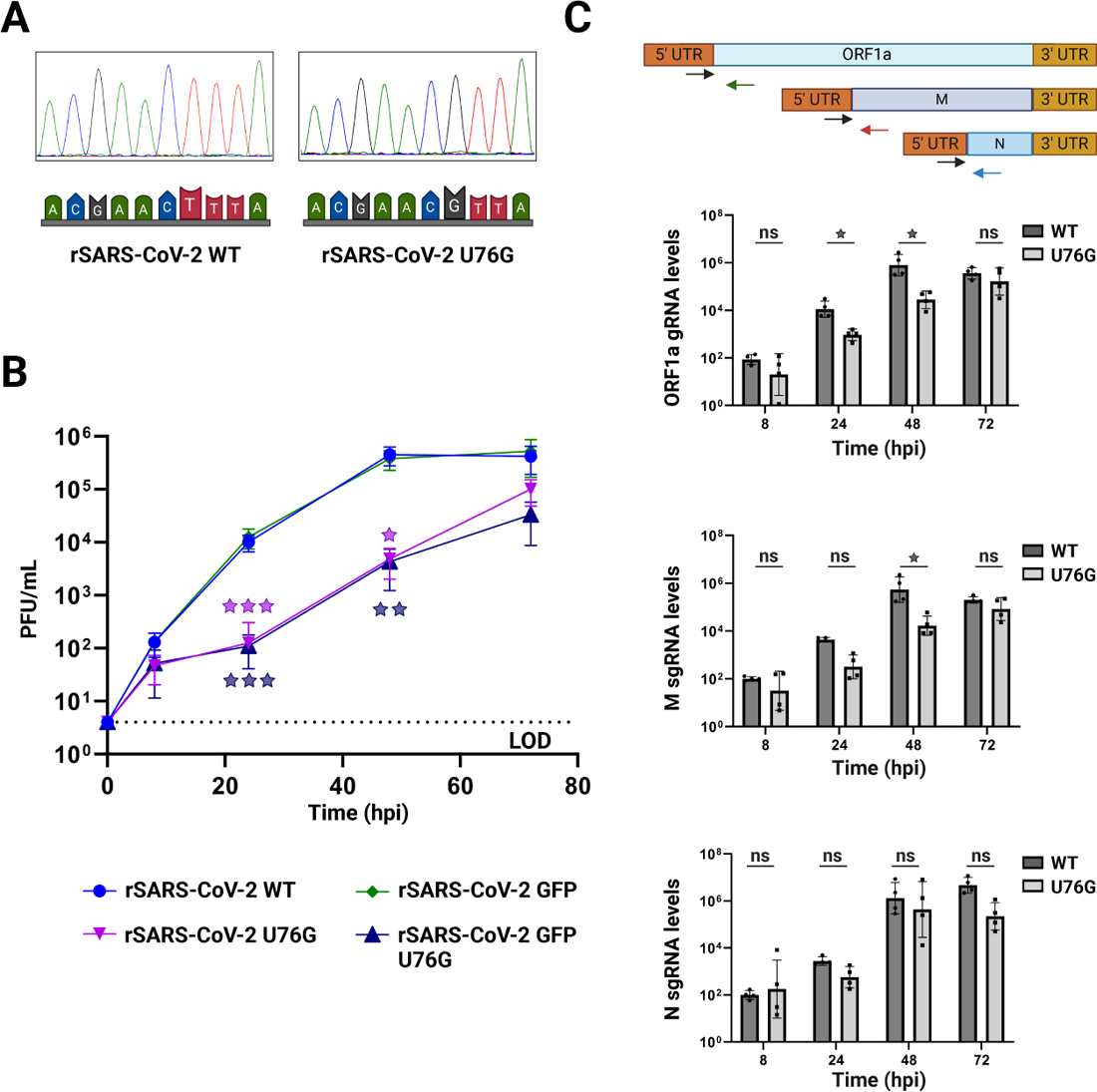
Characterization of SARS-CoV-2 5’UTR mutant. **(A)** Sanger sequencing of SARS-CoV-2 T76G mutant. Supernatant of rSARS-CoV-2 U76G, rSARS-CoV-2 GFP U76G and rSARS-CoV-2 WT infected cells were subjected to Sanger sequencing, showing the desired mutations compared to rSARS CoV-2 WT. **(B)** Growth kinetics of rSARS-CoV-2. Comparison of growth kinetics of rSARS-CoV-2 U76G and rSARS-CoV-2 GFP U76G mutant against rSARS-CoV-2 WT and rSARS-CoV-2 GFP in Vero E6 TMPRSS2 cells infected at a MOI 0.01 over a 3-day time course. *n* = 2 independent experiments with two replicates each. The graphic shows mean values ± SD. Stars indicate the degree of significance compared to the rSARS-CoV-2 WT or rSARS-CoV-2 GFP control (*=p<0.05, **= p<0.01, ***= p<0.001, ****= p<0.0001 by two-sided unpaired Student’s *t*-test). **(C)** gRNA and sgRNA expression. *Upper.* Schematic primer design to quantify gRNA and sgRNA expression. *Lower.* RT-qPCR of *ORF1a, M* and *N* RNA levels at 8, 24, 48 and 72 hpi in Vero E6 TMPRSS2 cells infected with rSARS-CoV2 WT or rSARS-CoV-2 U76G. Quantification relative to 18S rRNA. *n* = 2 independent experiments with two replicates each. The graphic shows mean values ± SD. Stars near the bars indicate the degree of significance compared to the rSARS-CoV-2 WT control condition (*=p<0.05, **= p<0.01, ***= p<0.001, ****= p<0.0001 by two-way ANOVA with Šidák’s test).

We then evaluated the replication kinetics of rSARS-CoV-2 U76G virus alongside WT. We used a MOI of 0.01 of both viruses to infect Vero E6 TMPRSS2 cells and collected samples at 8, 24, 48 and 72 hpi (**Figure 6B**). rSARS-CoV-2 U76G showed a striking growth deficiency at 24 and 48 hpi compared to rSARS-CoV-2 WT. Similar results were seen for rSARS-CoV-2 GPF U76G compared to rSARS-CoV-2 GFP, although at 72 hpi both viruses reached similar titers as their controls. Subsequently, we wondered whether the mutation U76G next to the TRS-L could impair the synthesis of sgRNAs leading to the reduced titers seen in replication kinetic assays. We used RT-qPCR primers encompassing the leader-body junction for *ORF1a* and also *M* and *N* sgRNA junctions^70^. We found significantly reduced amounts of sgRNA M at 48 hpi and gRNA ORF1a at 24 and 48 hpi. Meanwhile, the amount of sgRNA N was also reduced, although this was not statistically significant at any point evaluated (**Figure 6C**).

Overall, the incorporation of a U76G mutation adjacent to the TRS-L revealed a diminished viral growth kinetics, with reduced plaque size, and significantly reduced levels of sgRNA and gRNA compared to the wild-type virus.

## Discussion

Reverse genetics allows the engineering of recombinant and heavily modified RNA viruses, to reveal biological properties and better understand emerging viral pathogens. Here, we established a novel reverse genetics system for the rescue of recombinant SARS-CoV-2 virus. We achieved this by dividing and cloning the SARS-CoV-2 genome into six plasmids for propagation in *E. coli*, which is the lowest number of fragments reported to date^17,41,42^. Previously, the cloning and storage of long SARS-CoV-2 sequences in bacterial plasmids has been reported as unstable, which we have overcome by using low copy vectors. Whilst recombinant viruses can be produced directly from reverse transcribed RNA without cloning^41,42^, reverse transcription and amplification of the entire SARS-CoV-2 genome has not proven feasible meaning that genome segmentation and fragment assembly is still required. For these systems, the introduction of a desired mutation involves the generation of a new fragment, which may cause subsequent issues during the assembly of the whole genome. Using our system, mutations are introduced into plasmids without altering the number of fragments for the final assembly, and complex mutants can be generated more easily by combining fragments containing different mutations. Bacterial plasmids also have the advantage of being a permanent and easy to produce store of genetic material. They are also relatively easy to manipulate using established molecular techniques.

Several strategies have been previously described to assemble DNA fragments into a full SARS-CoV-2 genome, such as golden gate-like assembly using type II restriction enzymes^22,38^, in-yeast assembly^18^, bacterial artificial chromosome^23^ or polymerase extension reaction^41,42^. In our hands, fragment assembly using type IIs restriction enzymes did not provide the desired results due to incomplete assembly of the viral genome leading to a reduced amount of complete genome to be transfected into mammalian cells. On the other hand, polymerase extension, which involves the alignment of fragments containing 20 bp overlaps with the adjoining fragment, led to a higher assembly of the full viral genome, whilst also being shorter and more straightforward. As reported by others^24,41,42,44,47^, the inclusion of a linker sequence containing a CMV promoter, a HDV ribozyme to generate the authentic viral 3′ end containing a polyA tail, and SV40 polyA signal to ensure termination, allowed the assembly to be directly transfected into mammalian cells. Notably, this approach avoids expensive and complex RNA *in-vitro* production and RNA transfection^18,22,38^. Finally, we showed that PEI, which is a very low-cost transfection reagent, allowed the rescue of high titers of recombinant SARS-CoV-2 even after only one passage. Nanopore sequencing revealed high fidelity rescue (∼99.99% nucleotide accuracy, 99.97% amino acid accuracy after 2 passages), although there was some evidence for genome evolution during passaging in cell culture as reported before for other systems^41,42,73^.

We then used our reverse genetics system to explore the impact of different mutations on SARS-CoV-2 infection. We first focused on the receptor binding domain (RBD) of the spike protein due to its ongoing and rapid evolution since it’s a zoonotic transmission^7,48,49^. The mutations Y453F and N501Y emerged early in the pandemic and are reported to improve the binding of spike to receptor ACE2^7,52,5^>. Y453F was first identified in humans who had contact with infected minks^7^ and is located in a region of RBD interacting directly with ACE2^50,51^. This mutation has been associated with increased binding of the isolated RBD to hACE2 in biophysical^50^ and deep mutational screening assays^53^. In contrast, our results showed a significant decrease in binding of rSARS-CoV-2 Y453F to Vero E6 TMPRSS2 and A549 ACE2 overexpressing cells compared to rSARS-CoV-2 WT. It also showed slightly slower replication kinetics and reduced fitness during competition assays to rSARS-CoV-2 WT. In agreement with our results, Y453F mutation mostly enhances binding to mink ACE2^74^ and attenuates SARS-CoV-2 replication in human cells^51,75^. Together with the fact that Y453F has disappeared from human circulating SARS-CoV-2 strains, our data supports the notion that it is a Mink specific adaptation that has ‘spilled back’ to humans.

The second mutation N501Y has been widely investigated due to its early emergence and its continuing presence in Alpha, Beta, Gamma and Omicron variants of concern^7,48,49^. Most studies are in agreement that N501Y’s enhanced fitness is due to an increased affinity of the RBD for the human ACE2^52–5>^. Although N501Y may also reduce neutralization by certain RBD specific antibodies^7,76,77^. Here, we saw similar replication kinetics and cell binding of SARS-CoV-2 N501Y compared to rSARS-CoV-2 WT. Nevertheless, rSARS-CoV-2 N501Y outcompeted rSARS-CoV-2 WT over a period of 2 days in VERO E6 TMPRSS2 cells. Surprisingly, the fitness of rSARS-CoV-2 N501Y against WT was marginally reduced in A549 cells overexpressing ACE2. Whilst seemingly contradictory, our results could indicate that 501Y improves post-binding events. VERO E6 expresses both ACE2 and TMPRSS2^78^, in contrast to our A549 cell line only which only overexpress ACE2^62^. SARS-CoV-2 binds initially to ACE2 using Spike, followed by a process called “priming” where TMPRSS2 cleaves the spike protein at a specific site, known as the S2’ site^60,78^. This cleavage is essential for the fusion of the viral and cellular membranes and allows the viral membrane to merge with the host cell membrane to release the viral genetic material into the host cell^59,60^. Even if the binding of N501Y spike with ACE2 is not improved, post-binding events such as priming by TMPRSS2, membrane fusion or entry could be enhanced by N501Y mutation. Most studies reporting increased binding of N501Y with human ACE2 have been performed by computational modeling or *in vitro* studies using only the RBD domain and not in the context of full SARS-CoV-2 infection^54–57^. While two independent studies employed pseudotyped virus particles, the first found no difference in infectivity compared to WT during infection of VERO cells^79^, whereas the second reported only a small increase in infectivity of HIV-1 N501Y in HEK ACE2 cells^51^.

Furthermore, although N501Y virus was shown to replicate better than WT *in vivo* (in hamsters) or in human airway epithelial (HAE) cells, in Vero E6 cells the fitness difference was not as considerable^52^. This could be explained due to differences in the levels of ACE2 expression on the cell surface in these distinct systems. Taken together, our data highlights the complexity of investigating the relative fitness of spike mutants, and it becomes evident that various approaches and techniques need to be combined to enable meaningful evaluation of viral fitness. Biochemical, biophysical and cell culture-based assays using only the RBD are of fundamental importance but may fail to capture the effects of mutations on ACE2 affinity in the context of the full spike protein, as exemplified by the strong association with Y453F and increased binding of RBD to hACE2, which was not demonstrated in our rSARS-CoV-2 binding assay. Even when mutants are assessed in the context of the authentic virus, mutational studies should also consider the effects of epistasis which is known to be a powerful force affecting the evolution of protein sequences^80–82^. In particular, N501Y is known to shift the epistatic landscape of SARS-CoV-2^5^>. In our study, it must be noted that the N501Y mutant was assessed in the context of the ancestral Wuhan strain, which does not contain the D614G mutant that rapidly became dominant in circulating human strains^22,83,84^.

Finally, we explored a mutation adjacent to the TRS-L in the 5’-UTR of SARS-CoV-2. This single uridine residue, located at position 29,816 of negative sense RNA, was shown to be bound by NSP9^70^. NSP9 has capping and priming functions and interacts with the RdRp complex^67,68,71,72,70,71,85,86^. Our reverse genetics system allowed us to rescue and characterize rSARS-CoV-2 U76G, which leads to an exchange of Adenine for a Cytosine in the negative sense RNA. rSARS-CoV-2 U76G exhibited slower replication kinetics compared to WT at 24 and 48 hpi, with no noticeable difference at 72 hpi, the latest time point evaluated. Additionally, reduced levels of M sgRNA at 48 hpi and ORF1a gRNA at 24 and 48 hpi were observed when compared to WT. These findings suggest that the disruption of this site, located next to the TRS-L, impacts template switching or discontinuous transcription of positive-sense RNA, leading to decreased amounts of gRNA, sgRNA, and viral particles. However, multiple rounds of infections at later time, allows the sufficient accumulation of viral RNA and recovery of viral titers compared to rSARS-CoV-2 WT. While these results support the previously mentioned priming function of NSP9, it is crucial to acknowledge that we did not assess the binding of NSP9 to rSARS-CoV-2 U76G. Consequently, we cannot determine whether the observed phenotype is a consequence of reduced NSP9 binding or an unknown mechanism. Moreover, there are no prior reports detailing the role of this mutation, including its interaction with NSP9, nor other viral or cellular proteins. Further investigation is needed to fully understand its potential significance.

In conclusion, we have established a reverse genetics system that allows efficient, low-cost and flexible manipulation of the large genome of SAR-CoV-2. The use of recombinant SARS-CoV-2 viruses has the great advantage that they incorporate the effects of other viral or cellular processes during SARS-CoV-2 infection. The system was successfully used to generate and characterize two spike mutations seen in variants of concern, and a recently described NSP9 priming site in the 5’UTR revealing important effects on the viral life cycle. Our system therefore provides a robust and useful tool to rescue rSARS-CoV-2, which will not only to help the understanding of molecular and biological mechanisms involve in the infection and pathogenesis of SARS-CoV-2, but also potentially applicable to vaccine development or identification of therapeutic targets. We expect that the use of bacterial plasmids compatible with standard mutagenetic protocols, PCR based assembly and direct transfection with low-cost reagents will enable more widespread use of reverse genetics systems to study SARS-CoV-2.

## Methods

### Cells

The African green monkey kidney Vero E6 TMPRSS2, Human embryonic kidney HEK293T, HEK293 ACE2 and lung carcinoma epithelial cells A549 ACE2 were cultivated with Dulbecco’s Modified Eagle Medium (DMEM; Gibco) with 10% fetal bovine serum (Sigma-Alrich) and 100U/mL of Penicillin-Streptomycin (Gibco) under 5% CO_2_ and 37°C conditions.

### Plasmids

The full genome of SARS-CoV-2 WT was split into 6 fragments which were amplified using as a template the clone pCC1BAC-HIS3-SARS-CoV-2 and for SARS-CoV-2 GFP an extra fragment using as a template the clone pCC1BAC-HIS3-SARS-CoV-2-GFP both clones were kindly provided by Dr. Volker Thiel. The amplification was performed using high-fidelity PrimeSTAR GXL DNA polymerase (Takara). Next, fragments F1, F2, F3, F5, F6 WT and F6 GFP were cloned into the vector pUA66 and fragment F4 into vector Puc19.

The following plasmids were then modified: for plasmid F1 the initial promoter T7 was removed and replaced by a CMV promoter. In the case of fragment F6 WT and F6 GFP a linker sequence was added to the 3’UTR. The linker contains the next elements: 1) Hepatitis delta virus ribozyme (HDVr), 2) SV40 poly (A) signal and 3) a spacer sequence of 364bp.

Mutations N501Y, Y453F and T76G were inserted in plasmid F5 or F1 respectively by Directed Site Mutagenesis using the primers (**Supplementary Table 2**) and high-fidelity PrimeSTAR GXL DNA polymerase (Takara).

All plasmids were subjected to Sanger sequencing and only a single silent point mutation at position 28,103 located in Fragment 6 WT was detected compared to the sequence of SARS-CoV-2 isolate Wuhan-Hu-1 (GenBank MN996528.3).

An individual plasmid containing the nucleoprotein (N) of SARS-CoV-2 was generated using the clone pCC1BAC-HIS3-SARS-CoV-2 as a template. The N sequence of 1260 bp was amplified using the primers (**Supplementary Table 2**) and a CMV promoter added upstream of the sequence. Both fragments were cloned into the vector Puc19. Analysis by Sanger sequencing did not show any change in the sequence.

### PCR amplification of SARS-CoV-2 individual fragments

Each individual SARS-CoV-2 fragment was amplified from their respective plasmids using an exclusive pair of primers (**Supplementary Table 2**) and high-fidelity PrimeSTAR GXL DNA polymerase (Takara), followed by gel isolation with NucleoSpin Gel and PCR Clean-up (Macherey-Nagel) following the manufacturer’s recommendations. PCR conditions consisted in 0.05 U of PrimeSTAR GXL polymerase (Takara Bioscience), 250 nM of each primer, 200 µM of each dNTP and x1 PrimerSTAR GXL buffer in a total volume of 50 µL. Cycling conditions were initial denaturation for 2 min at 98°C, followed by 35 cycles for 10 sec at 98°C, 15 sec at 55°C, and 10 min at 68°C, followed by a final extension for 15 min at 68°C. Amplicon quality was checked on 1% agarose gel post-stained in EtBr.

### SARS-CoV-2 genome assembly

Recombinant SARS-CoV-2 was assembled by CPER, each fragment possesses complementary ends of 20 nucleotides overlap used for the assembly. Six fragments were mixed in equimolar amount of 0.1 pM, 2 µL of PrimeSTAR GXL DNA polymerase (Takara), 200 uM of each dNTP, 1x GXL buffer into a final volume of 50 µL.

Two initial cycling conditions were tested: 1) initial denaturation at 98°C for 30 sec, 12 cycles of 10 sec at 98°C, 20 sec at 55°C and 10 min at 68°C, and a final elongation for 12 min at 68°C. 2) initial denaturation at 98°C for 2 min, 20 cycles of 10 sec at 98°C, 15 sec at 55°C and 25 min at 68°C, and a final elongation for 25 min at 68°C.

The first condition was chosen due to the generation of a band with the desired size (30 kb). We performed an optimization of the number of cycles using 6, 9 or 12 cycles. Once again, the condition using 12 cycles performed the best results and all the recombinant SARS-CoV-2 viruses were generated using the same conditions but exchanging the corresponding plasmid with the desired mutation. Assembly reactions were then used for transfection without any purification step.

### Genome assembly transfection and virus rescue

Virus rescue was accomplished by reverse transfection of SARS-CoV-2 assembly into HEK293T ACE2 cells. As an initial test different conditions were tested: Briefly, 25 µL of assembly without purification were mixed with 7.2 or 14.4 µL of polyethylenimine (PEI) and 50 µL of assembly without purification were mixed with 14.4 or 28.8 µL of PEI, the mixture was placed in a 6 well plate/dish. After 10 min of incubation at room temperature HEK293T ACE2 were dropped on the top of the DNA: PEI mixture. Cells were incubated under 5% CO_2_ and 37°C conditions. After 24 h, cells were trypsinized and seeded over a monolayer of Vero E6 TMPRSS2 cells, supernatants were collected after 10 days. Recombinant viruses were amplified once on Vero E6 TMPRSS2 in order to generate viral stocks. For the following experiments the only condition used was the reverse transfection of 50 µL of assembly mixed with 28.8 µL of PEI.

### RNA extraction, cDNA synthesis and qPCR

Total RNA was extracted with Trizol (Invitrogen) using the manufacturer’s recommendations and the totality of the viral RNA was treated with Turbo DNase (Thermo Fisher Scientific) for 30 min at 37°C. Following DNase treatment, RNA was column purified using NTC buffer and the NucleoSpin Gel and PCR Clean-up kit (Macherey-Nagel), according to the manufacturer’s instructions.

The RNA was then reverse transcribed using SSIV (Thermo Fisher Scientific) with a set of random hexamers (IDT). Quantitative real-time PCR (qRT-PCR) was performed using PowerUP SYBR green (Thermo Fisher Scientific) according to manufacturer’s instructions. A standard curve for the Nucleocapsid (*N*) and RNA polymerase dependent of RNA (*RdRp*) was made using serial dilutions of plasmids F6 and F4 respectively. The oligonucleotides used to amplify Nucleocapsid (*N*), RNA polymerase dependent of RNA (*RdRp*) and 18s rRNA are described in **Supplementary Table 2**. The level of each RNA was determined by CFX96 Touch Real-Time PCR Detection System (Bio-Rad) with the cycling condition: 50°C for 2 min, 95°C for 2 min, followed by 40 cycles of 95°C for 15 s and 60°C for 30 s, finishing with melt profile analysis. The software used for data statistical analysis is Prism 7 (GraphPad).

### cDNA synthesis and Sanger sequencing for spike mutants and 5’UTR mutant

After RNA extraction and DNAse treatment, the RNA was then reverse transcribed for 4 h using SSIV (Thermo Fisher Scientific) with a set of reverse SARS-CoV-2 primers (**Supplementary Table 2**). An amplicon of 2.4 kb was produced using the previous cDNA and a specific pair of primers for the spike mutants Y453F and N501Y (Fw primer: ACAAATCCAATTCAGTTGTCTTCCTATTC and Rv primer: TGTGTACAAAAACTGCCATATTGCA) or for T76G mutant (Fw primer: ACCAACCAACTTTCGATCTCTTGT and Rv primer: GCTTCAACAGCTTCACTAGTAGGT) using the following conditions: 1 µL of diluted 1/10 RT reaction with 0.05 µL of PrimeSTAR GXL polymerase (Takara Bioscience), 250 nM of each primer, 200 µM of each dNTP and x1 PrimerSTAR GXL buffer in a total volume of 25 µL. Cycling conditions were initial denaturation for 1.5 min at 98°C, followed by 35 cycles for 10 sec at 98°C, 15 sec at 55°C, and 3 min at 68°C, with a final extension for 5 min at 68°C. Amplicon quality was checked on 1 % agarose gel post-stained in EtBr.

PCR products were purified using NucleoSpin Gel and PCR Clean-up (Macherey-Nagel) according to manufacturer’s recommendations. Spike mutant PCR products for Y453F and N501Y were analyzed by Sanger sequencing using the specific primer (TTCAGCCCCTATTAAACAGCCTGCACGTGT) meanwhile PCR product for T76G was analyzed by Sanger sequencing with the specific primer (GGCAAAACGCCTTTTTCAACTTC).

### Viral infections and growth kinetics

In general, rSARS-CoV-2 inoculums were prepared in DMEM supplemented with 1% FCS. Before the infection, Vero E6 TMPRSS2 cells were washed once with PBS and incubated with the respective inoculum for 1 h at 37°C with gently shake every 10 min. The inoculum was removed and fresh DMEM supplemented with 5% FCS, 100 mg/mL of penicillin and 100 U/mL of streptomycin was added to the cells.

Replication kinetics for the different recombinant SARS-CoV-2 were performed using Vero E6 TMPRSS2 cells with a MOI 0.01. Samples collected at 8, 24, 28 and 72 hpi were titrated in duplicate by plaque assay. Statistical analysis comparison was performed using two-sided unpaired Student’s t-test with Prism 7 (GraphPad).

### Plaque assay

Cell supernatant containing virus were 10-fold serial diluted in DMEM 1% FCS, inoculated onto TMPRSS2-Vero E6 cell monolayer in duplicate, incubated at 37°C for 1 h. After the incubation, the inoculum was removed and the cell monolayer was overlayed with 0.6% (w/v) methylcellulose (Carl Roth) in MEM (Gibco) supplemented with 25 mM of HEPES, 0.44% NaHCO3, 2 mM of GlutaMAX (Gibco), 100 U mL−1 of streptomycin, 100 mg mL−1 of penicillin and 5% FCS and incubated at 37°C. After 3 days, cells were fixed and stained with 2x staining solution (0.23% crystal violet, 8% formaldehyde, 10% ethanol) directly to the medium for 24 h. Cells were washed twice with H_2_O and plaques enumerated to determine viral titers.

### Nanopore sequencing and bioinformatic analysis

Supernatant of infected VERO E6 TMPRRS2 cells were treated for RNA extraction using Trizol (Invitrogen) based on the manufacturer’s recommendations, followed by Ethanol/Sodium Acetate precipitation and resuspension in RNase-free H2O. Then, 10 ug of RNA were treated with Turbo DNase (ThermoFisher Scientific) for 30 min at 37°C. Following DNase treatment, RNA was column purified using the NucleoSpin Gel and PCR Clean-up kit (Macherey-Nagel) with NTC buffer according to the manufacturer’s instructions.

Reverse transcription was performed using MarathonRT. pET-6xHis-SUMO-MarathonRT encoding MarathonRT was a gift from Anna Pyle (Addgene plasmid # 109029; http://n2t.net/addgene:109029; RRID: Addgene_109029) and purified according to (PMID 29109157). Viral RNA was reverse transcribed using a mix of reverse primers listed in **Supplementary Table 2**. Specifically, RNA was mixed with 0.5 mM dNTPs, 5 uM of each primer in 9 µL total volume and denatured for 5 min at 65°C. Samples were placed on ice for 2 min and reverse transcription was initiated by adding 40 U of MarathonRT in 50 mM Tris-HCl pH 8.3, 200 mM KCl, 20% glycerol (v/v), 1 mM MnCl_2_, 4 U of RNasin in a 20 µL total volume. Samples were incubated for 4 h at 42°C. Controls lacking reverse transcriptase were carried out as above, with the omission of the MarathonRT enzyme.

The RT product was subsequently used in a set of 14 separate PCR reactions to produce 2.4 kb amplicons that covered the full genome of SARS-CoV-2. PCR amplification conditions were 1 µL of diluted 1/10 RT reaction with 0.05 U of PrimeSTAR GXL polymerase (Takara Bioscience), 250 nM of each primer, 200 µM of each dNTP and x1 PrimerSTAR GXL buffer in a total volume of 25 µL. Cycling conditions were initial denaturation for 1.5 min at 98°C, followed by 35 cycles for 10 sec at 98°C, 15 sec at 55°C, and 3 min at 68°C, followed by a final extension for 5 min at 68 °. Amplicon quality was checked on 1 % agarose gel post-stained in EtBr.

PCR products were pooled and purified via SPRI bead purification (Mag-Bind Totalpure NGS, Omega Bio-tek) by addition of 0.6 x volumes of beads followed by light agitation for 5 minutes at room temperature. Beads were pelleted on a magnetic rack (Invitrogen DYNAL), followed by removal of supernatant and 2 washes with 100 µL freshly prepared 70% ethanol. Finally, beads were air dried for 3-5 min (until appearance changed from glossy to rough) and DNA was eluted by addition of 30 µL H_2_O, followed by 5 min incubation at room temperature.

DNA concentration was quantified via Nanodrop, and 300 ng of pooled product was taken into the Nanopore native ligation barcoding library preparation. Specifically, the DNA in 11.5 µL in a PCR tube was end-repaired by addition of 1.75 µL NEB Ultra II End Repair Buffer (NEB) and 0.75 µL Enzyme Mix followed by thorough mixing via pipetting, incubated for 5 min at room temperature and 5 min at 65°C. Next, 1 µL of end-repaired DNA was transferred into a new PCR tube, followed by addition of 0.75 µL H_2_O, 1.25 µL native ligation barcode (ONT SQK-NBD114-96) and 3 µL Blunt/TA Ligase Master Mix (NEB). The reaction was mixed by pipetting, incubated for 20 min at room temperature, and terminated by addition of 1 µL EDTA (SQK-NBD114-96). The barcoded DNA samples were then pooled and purified by addition of 0.4 volumes of SPRI beads (SQK-NBD114-96). For binding, they were incubated for 5 min with mixing, then pelleted on a magnetic rack (DYNAmag). Next, beads were washed twice by resuspension and re-pelleting in 200 µL Short Fragment Buffer (SFB, ONT SQK-NBD114-96) before a final wash step with 100 µL 80 % ethanol. After a brief drying on air, the DNA was then eluted by addition of 35 µL and incubation for 10 min at 37°C with mixing. Finally, 30 µL of the barcoded DNA was ligated onto 2.5 µL Native Ligation Adapter (NA, ONT SQK-NBD114-96) by addition of 12.5 µL NEBNext Quick Ligation Reaction Buffer (NEB) and 5 µL high conc. T4 DNA Ligase (NEB). After incubation for 20 min at room temperature, 20 µL of Ampure XP Beads (SQK-NBD114-96) were added, followed by incubation for 10 min at room temperature with mixing, pelleting of beads in a magnetic rack, and two washes with 125 µL SFB (SQK-NBD114-96). After removal of wash buffer, the beads were resuspended in 7 µL Elution Buffer (SQK-NBD114-96), incubated for 10 min at 37°C, followed by pelleting and transfer of the supernatant into a 1.5 mL DNA LoBind tube. The prepared library was quantified with AccuClear Ultra High Sensitivity dsDNA kit and 14 ng (approximately 10 fmol) were loaded onto a Kit 14 Flongle flow cell (FLO-FLG114). Sequencing data was acquired with MinKNOW version 22.12.7 (4 kHz sampling) for the rescue verification and 23.07.8 (5 kHz sampling) for the competition assay respectively.

All data was subsequently basecalled and demultiplexed with dorado v0.5.0 (min-qscore 11) and aligned to the respective SARS-CoV-2 wildtype (NC045512.2) or SARS-CoV-2 GFP (BioProject: PRJNA615319; BioSample: SAMN14450690) reference sequence with LAST v1450. The generated maf files were then converted into sorted and indexed bam files using samtools v1.16.1, and mutation frequencies for each position with a coverage of at least 100 were quantified with perbase v0.8.5. The generated tables were then parsed into pandas (v1.4.3) dataframes, analyzed with custom python code provided as Jupyter notebooks in the supplementary data. Figures were generated with the python package plotly v5.9.0.

### Binding assay

Vero E6 TMPRSS2 and A549 ACE2 cells were washed twice with cold PBS and then incubated with rSARS-CoV-2 WT, rSARS-CoV-2 Y453F or rSARS-CoV-2 N501Y for 1 h at 4°C with gentle shaking every 10 min. Then, the inoculum was removed, and cells were washed twice with cold PBS and cells were lysed with TRIZOL. RNA extraction, RT and qPCR for 18S rRNA and *N* were performed as described before. To calculate differences in RNA expression we used the DDCT method versus 18S rRNA. Statistical analysis comparison between binding of each RDP spike mutant against rSARS-CoV-2 WT was performed using one-way ANOVA with Prism 7 (GraphPad).

### Competition assay

Competition assay was performed infecting a monolayer of VERO E6 TMPRSS2 or A549 ACE2 cells with 5 different virus combinations at a MOI 0.1: 1) rSARS-CoV-2 WT, 2) rSARS-CoV-2 Y453F, 3) rSARS-CoV-2 N501Y, 4) rSARS-CoV-2 WT plus rSARS-CoV-2 Y453F or rSARS-CoV-2 WT plus rSARS-CoV-2 N501Y for 1h at 4°C with gently shake every 10 min. After, the inoculum was removed, fresh DMEM supplemented with 5% FCS, 100 mg/mL of penicillin and 100 U/mL of streptomycin were added to the cells.

After 2 days, supernatant and cell monolayer were lysed using TRIZOL LS or TRIZOL respectively. RNA extraction and RT using Marathon RT were performed as described before. Next, each sample was PCR amplified to produce a 2.4 kb amplicon using primers (Fw primer: ACAAATCCAATTCAGTTGTCTTCCTATTC and Rv primer: TGTGTACAAAAACTGCCATATTGCA). Library preparation, nanopore sequencing and data analysis were carried out as described in the section for nanopore sequencing and bioinformatic analysis. In the case of VERO E6 TMPRSS2 cells two independent experiments were done meanwhile with A549 ACE2 only one independent experiment was performed.

### gRNA and sgRNA quantification

VERO E6 TMPRSS2 cells were infected with rSARS-CoV-2 WT or rSARS-CoV-2 at a MOI 0.1 collecting samples at 8, 24, 48 and 72 hpi. After RNA extraction and DNAse treatment described as before, RNA was reverse transcribed using SSIV (Thermo Fisher Scientific) with a set of random hexamers (IDT). To specifically analyze viral sgRNAs and gRNA, we used qRT-PCR with previously designed primers^70^ (**Supplementary Table 2**): a forward primer that binds within the SARS-CoV-2 leader sequence and a specific reverse primer for the ORF1a RNA, the M mRNA or the N mRNA. qRT-PCR was performed using PowerUP SYBR green (Thermo Fisher Scientific) according to manufacturer’s instructions. To calculate differences in RNA expression we used the DDCT method versus 18S rRNA. Statistical analysis comparison between rSARS-CoV-2 U76G against rSARS-CoV-2 WT at every time point was performed using two-way ANOVA with Prism 7 (GraphPad).

### Microscopy

Due to SARS-CoV-2 is considered as a BioSafety Level 3 (BSL3) pathogen, all the images of live cells provided in this studio were performed inside the BSL3 laboratory using a Revolve R4 microscope (ECHO).

## Acknowledgements

We would like to that Prof. Dr. Volker Thiel and Prof. Dr. Chase Beisel for providing the SARS-CoV-2 YAC and pUAGG plasmids, respectively. We would also like to thank Prof. Dr. Andreas Pichlmair and Prof. Dr. Stefan Pöhlmann for generating and providing the A549 ACE2 cells and VERO E6 TMPRSS2, respectively. We also thank Lukas Pekarek for his expertise and advice in cloning. We thank Dr. Christine Krempl, Uddhav Ambi, Shazeb Ahmad, Liqing Ye, Sebastian Zielinski for expert technical assistance and feedback. Funding for this study was provided by the Helmholtz Association (VH-NG-1347 to RS). MON received a postgraduate fellowship from the Helmholtz Institute for RNA-based Infection Research (HIRI) Graduate Programme. The funders had no role in study design, data collection and analysis, decision to publish or preparation of the manuscript.

## Author contributions

MON and RPS conceived the study. MON, LD, MM, RPS designed the experiments. MON, PB, TH, NS, CB, ASG performed the experiments. MON and RPS wrote the manuscript in consultation with the other authors.

## Data availability

Sequencing data will be deposited at ENAXXX upon publication.

---

**Supplementary Table 1.** List of mutations in SARS-CoV-2 sequencing with more 50% presence.

| Virus | Passage | Rescue | Position | Reference<br>base | Mutation | Transversion<br>(TV) or<br>Transition (TS) | %<br>Mutation<br>Presence | Amino<br>acid<br>change | Protein<br>altered |
| --- | --- | --- | --- | --- | --- | --- | --- | --- | --- |
| WT | 1 | 1 | 29662 | T | C | TS | 92.34 | 3' UTR | - |
| WT | 1 | 2 | 3426 | C | T | TS | 94.22 | P236L | NSP3 |
|  | 1 | 2 | 7299 | C | T | TS | 97.03 | A1527V | NSP3 |
|  | 1 | 2 | 9698 | A | G | TS | 98.41 | I382V | NSP4 |
|  | 1 | 2 | 28320 | C | T | TS | 95.59 | T17M | N |
| WT | 1 | 3 | 5173 | C | T | TS | 50.85 | Syn. | - |
|  | 1 | 3 | 5884 | C | T | TS | 52.10 | Syn. | - |
| WT | 1 | 4 | 3199 | T | C | TS | 55.66 | Syn. | - |
| WT | 2 | 1 | 6258 | C | G | TV | 97.35 | T1180S | NSP3 |
|  | 2 | 1 | 6633 | C | G | TV | 96.61 | A1305G | NSP3 |
|  | 2 | 1 | 23663 | G | A | TS | 96.40 | A701T | S |
|  | 2 | 1 | 28810 | C | T | TS | 96.35 | Syn. | - |
| GFP | 1 | 1 | 25962 | T | C | TS | 70.66 | Syn. | - |
| GFP | 2 | 1 | 5896 | A | T | TV | 86.82 | Syn. | - |

**Supplementary Table 2.**
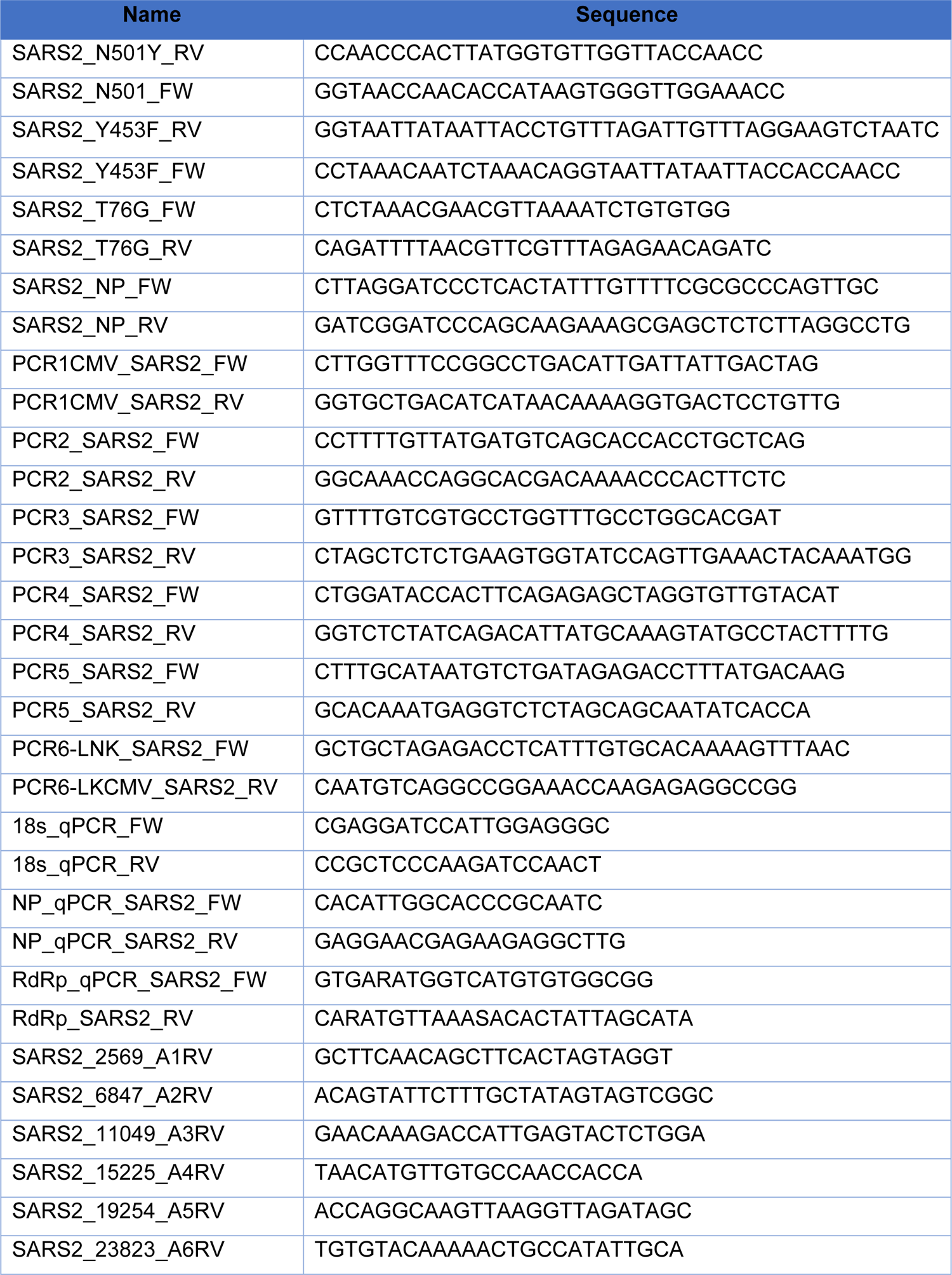
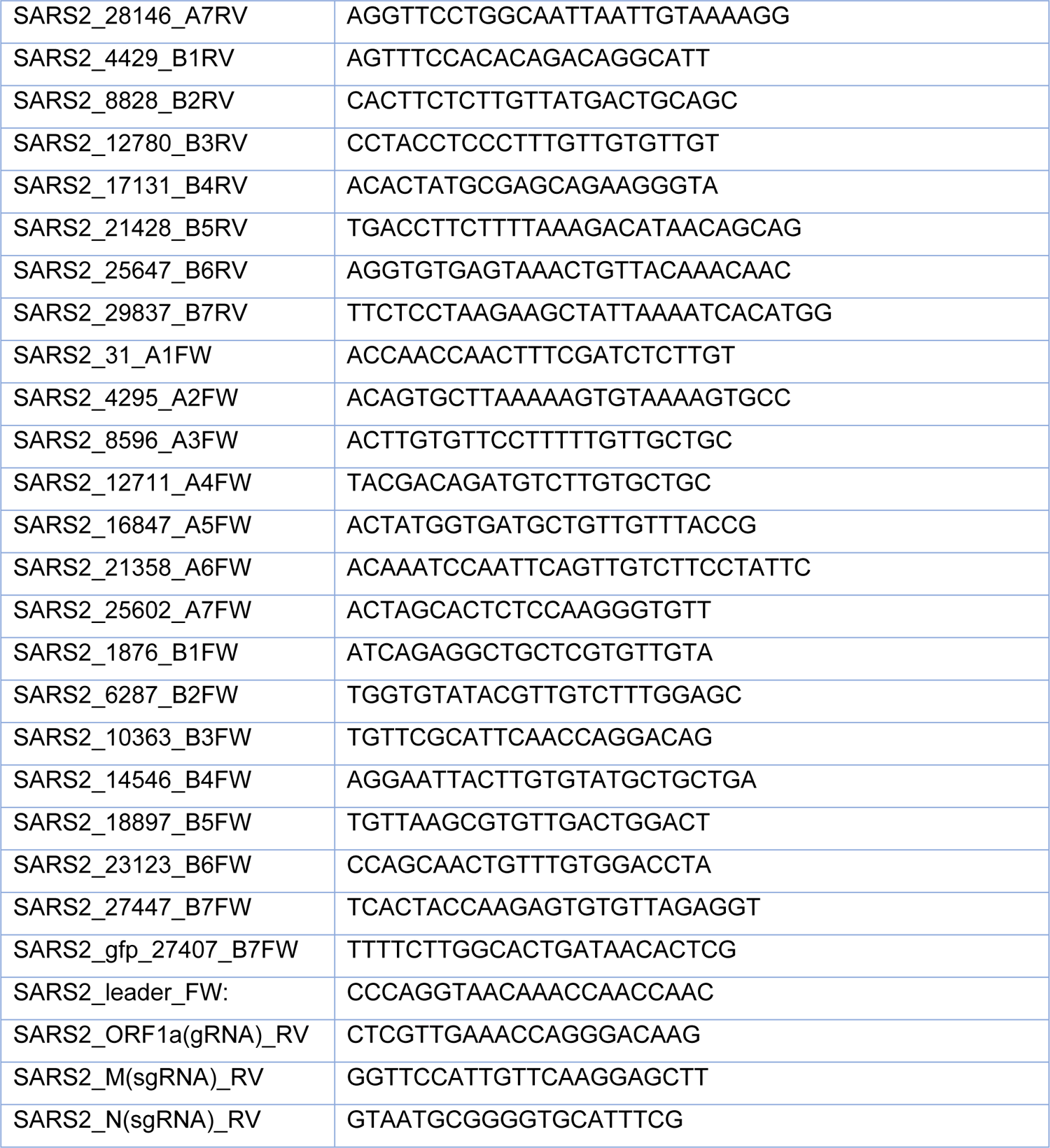
List of oligonucleotides.

**Supplementary Figure 1.**
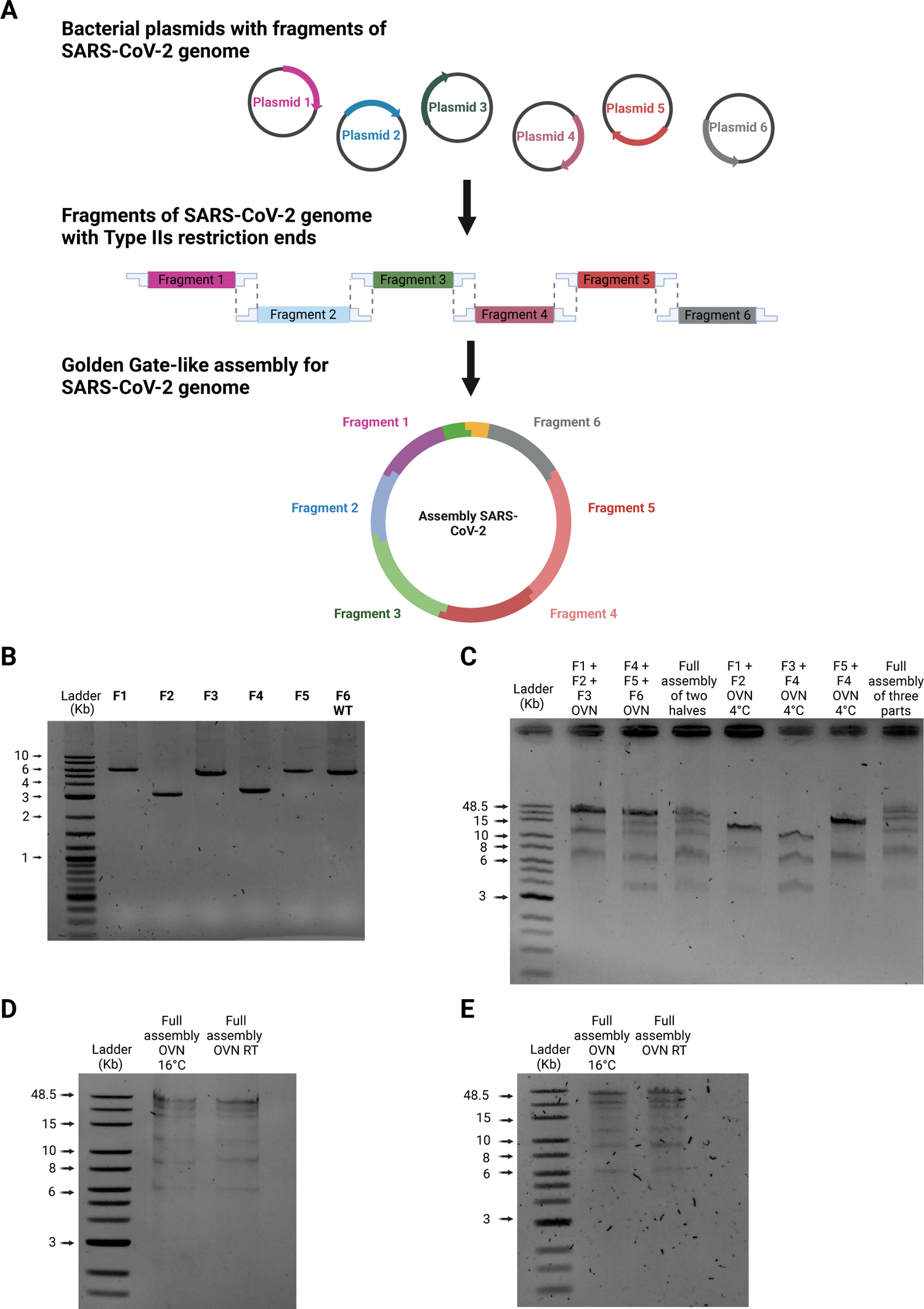
Reverse genetics system for SARS-CoV-2 using Golden Gate assembly. **(A)** SARS-CoV-2 genome is split into 6 fragments and cloned into bacterial plasmids. After *Bsa*I digestion, each fragment has complementary ends suitable for ligation and assembly of the complete viral genome. **(B)** DNA fragments of SARS-CoV-2 after plasmid digestion using *Bsa*I. **(C)** Partial assembly of DNA SARS-CoV-2. DNA fragments were mixed in two reactions containing F1+F2+F3 and F4+F5+F6 (lines 1 and 2) or in three reactions with F1+F2, F3+F4 and F5+F6 (lines 4, 5 and 6) incubated at 4°C overnight with T4 DNA ligase. After the incubation, the individual reactions were mixed for the two reactions (line 3) and the three reactions (line 7) and incubated again at 4°C overnight with T4 DNA ligase to assemble the full genome of SARS-CoV-2. **(D)** Full assembly of SARS-CoV-2. Two temperatures (16°C and RT) were tested overnight for the full assembly of SARS-CoV-2 genome when the partial assembly involved two reactions or **(E)** when the partial assembly involved three reactions.

**Supplementary Figure 2.**
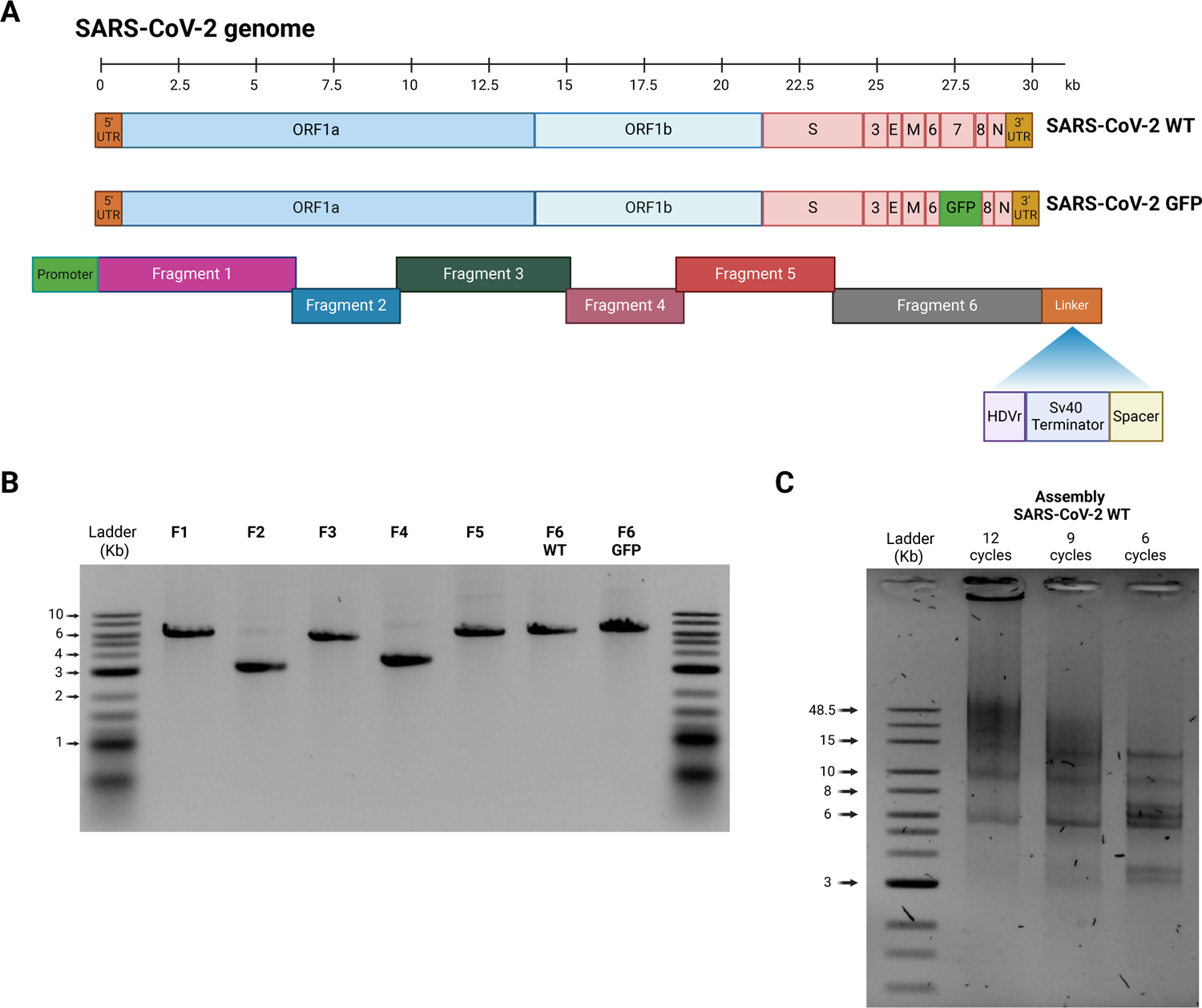
Design and optimization of polymerase extension reaction for SARS-CoV-2 DNA assembly. **(A)** Schematic representation of SARS-CoV-2 WT and rSARS-CoV-2 GFP. Some modifications were introduced into the SARS-CoV-2 genome. First, a CMV promoter was added to the upstream region of the 5’UTR in fragment F1. Next, a linker sequence was introduced after the 3’UTR in fragments F6 WT and F6 GFP consisting of: 1) *Hepatitis delta virus ribozyme*: cleavage of the phosphodiester bound for a precise 3’ end of the vRNA; 2) *SV40 Poly a terminator:* ending of transcription to avoid the generation of concatemeric RNA and 3) *a Spacer sequence:* 350 bp to create an intermediate area between Sv40 and the CMV transcription promoter during the circularization assembly. **(B)** DNA fragments of SARS-CoV-2. PCR products of 7 fragments of SARS-CoV-2, including the fragment for the GFP reporter, using bacterial plasmids as template. **(C)** Polymerase extension assembly for SARS-CoV-2. Three different assembly conditions (12, 9 and 6 polymerase extension cycles) were tested using an equimolar mixture of 6 DNA fragments to assemble the 30 kb SARS-CoV-2 genome.

**Supplementary Figure 3.**
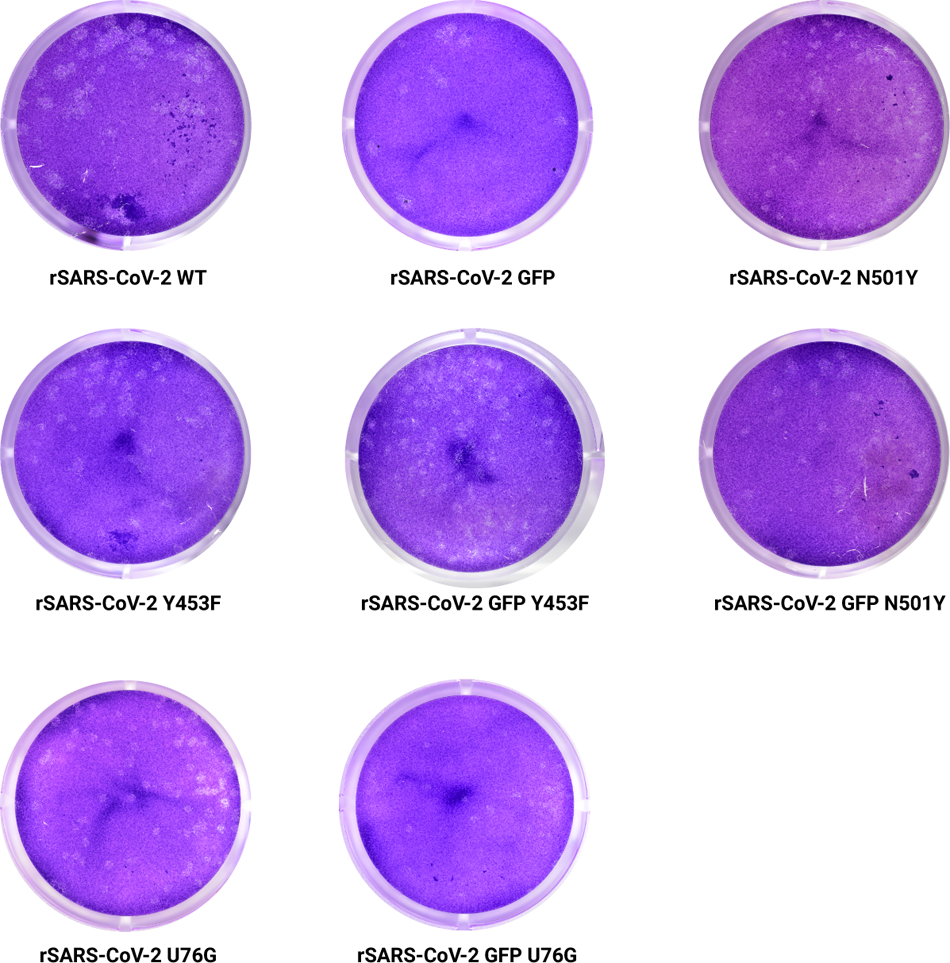
Plaque morphology of recombinant SARS-CoV-2. Representative plaque morphology of rSARS-CoV-2 WT, rSARS-CoV-2 GFP, rSARS-CoV-2 N501Y, rSARS-CoV-2 Y453F, rSARS-CoV-2 GFP Y453F, rSARS-CoV-2 GFP N501Y, rSARS-CoV-2 U76G and rSARS-CoV-2 GFP U76G in Vero E6 TMPRSS2 at 72 hpi.

**Supplementary Figure 4.**
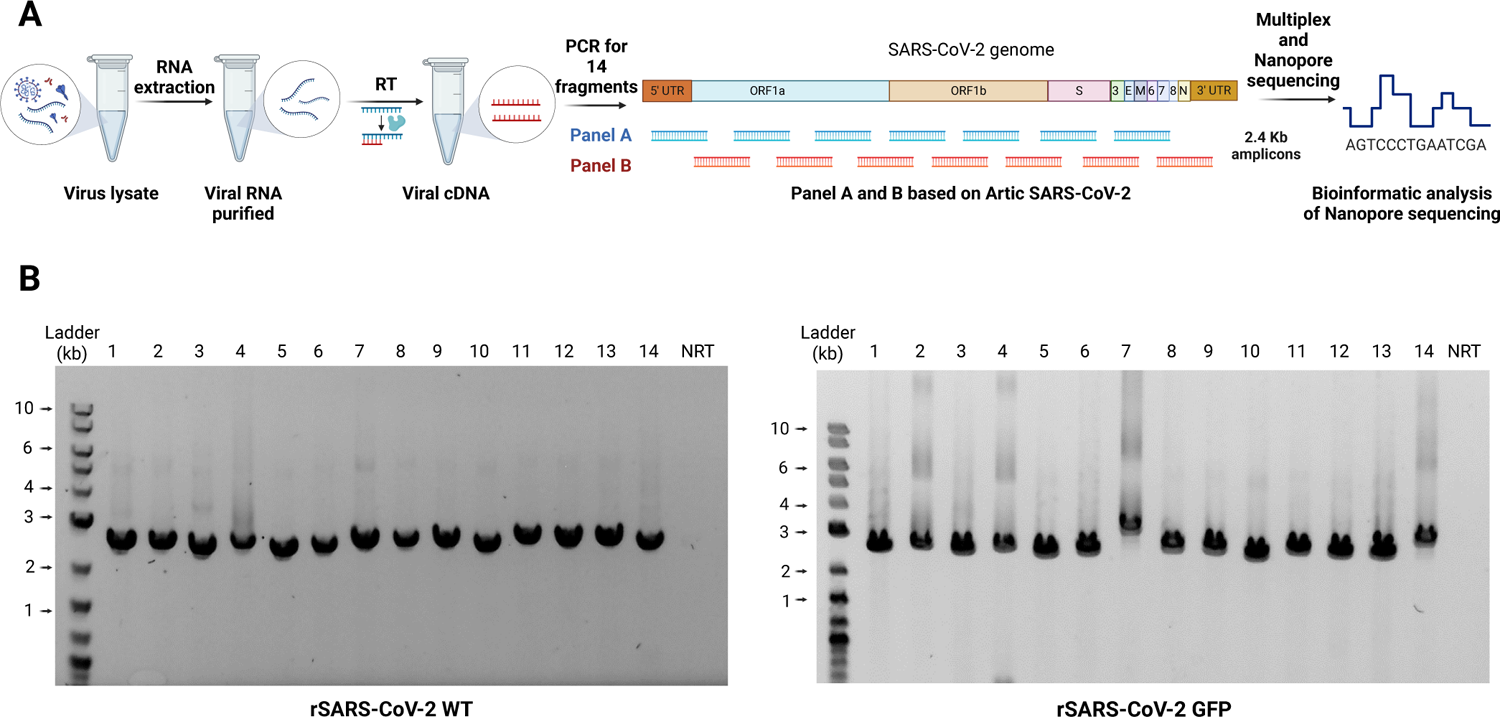
Design Oxford Nanopore sequencing for rSARS-CoV-2. **(A)** Oxford Nanopore DNA sequencing workflow for rSARS-CoV-2. **(B)** DNA fragments of SARS-CoV-2 for Nanopore sequencing. 14 amplicons of 2.4 kb covering the full genome of rSARS-CoV-2 WT *(left panel)* and rSARS-CoV-2 GFP *(right panel)* were generated by RT-PCR of RNA extracted from supernatant of Vero E6 TMPRSS2 cells at 72 hpi.

**Supplementary Figure 5.**
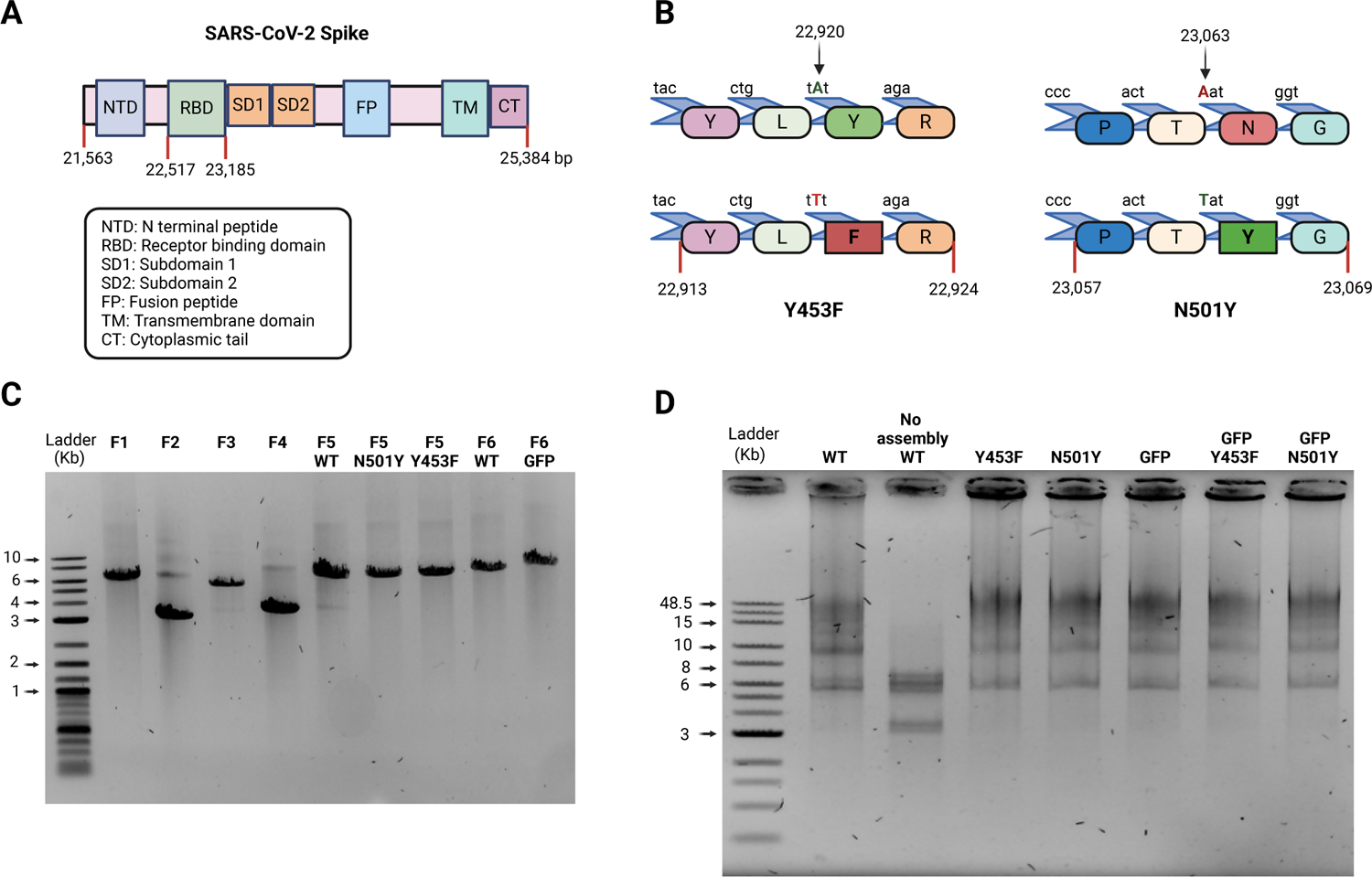
Generation of SARS-CoV-2 spike mutants. **(A)** Schematic representation of SARS-CoV-2 spike. Spike sequence is located in the second third of the SARS-CoV-2 genome with an extension of 3,801 bp producing a protein composed of 1,267aa. Spike possesses different domains including the RBD (Receptor Binding Domain). **(B)** Visual representation of RBD spike mutants. Mutation Y453F and N501Y mutations were introduced in fragment 5, at positions 23,063 and 22,920 of SARS-CoV-2 genome. In both mutations, a single nucleotide was changed in order to alter the amino acid sequence. **(C)** DNA fragments of SARS-CoV-2. PCR products of 9 fragments of SARS-CoV-2, including the fragment 5 with Y453F and N501Y mutation. **(D)** Assembly of complete DNA SARS-CoV-2. An equimolar mixture of 6 DNA fragments is assembled in the 30 kb SARS-CoV-2 genome, in order to generate different recombinant SARS-CoV-2. A control including all the fragments but without DNA polymerase was performed.

**Supplementary Figure 6.**
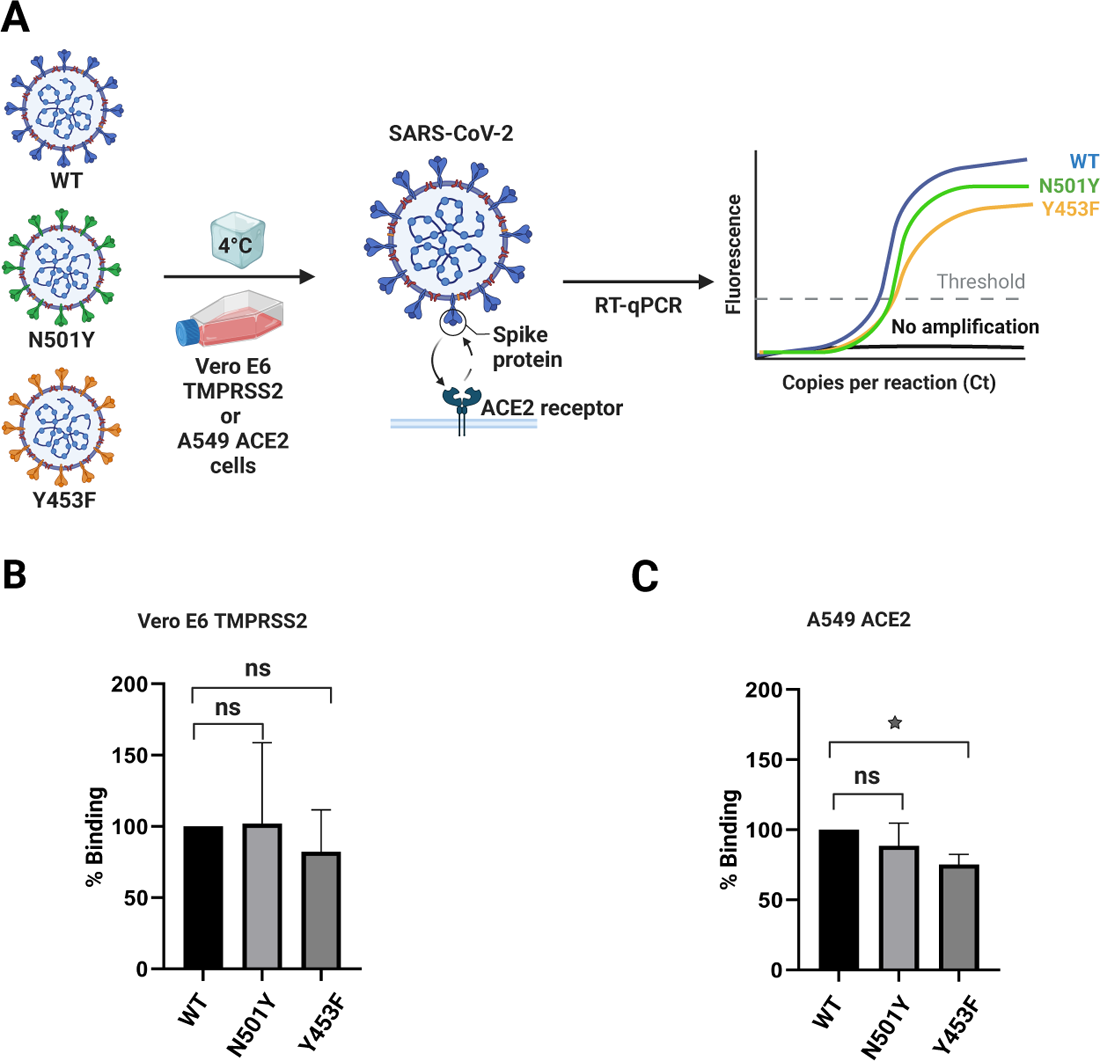
Binding assay for recombinant SARS-CoV-2 RBD spike mutants. **(A)** Schematic representation of Binding assay. **(B** and **C)**. Binding assay for SARS-CoV-2 spike mutants. Vero E6 TMPRSS2 cells or A549 ACE2 were incubated with rSARS-CoV-2, rSARS-CoV-2 Y453F or rSARS-CoV-2 N501Y at 4°C to evaluate the interaction between the spike and ACE2 receptor. Samples were evaluated by qPCR for *RdRp* and compared to rSARS-CoV-2 WT in **(B)** Vero E6 TMPRSS2 at a MOI 0.01 and **(C)** A549 ACE2 at a MOI 0.01. Quantification relative to 18S rRNA. *n* = 3 independent experiments with two replicates each. The graphic shows mean values ± SD. Stars above the bars indicate the degree of significance compared to the control condition (*=p<0.05, **= p<0.01, ***= p<0.001, ****= p<0.0001 by one-way ANOVA).

**Supplementary Figure 7.**
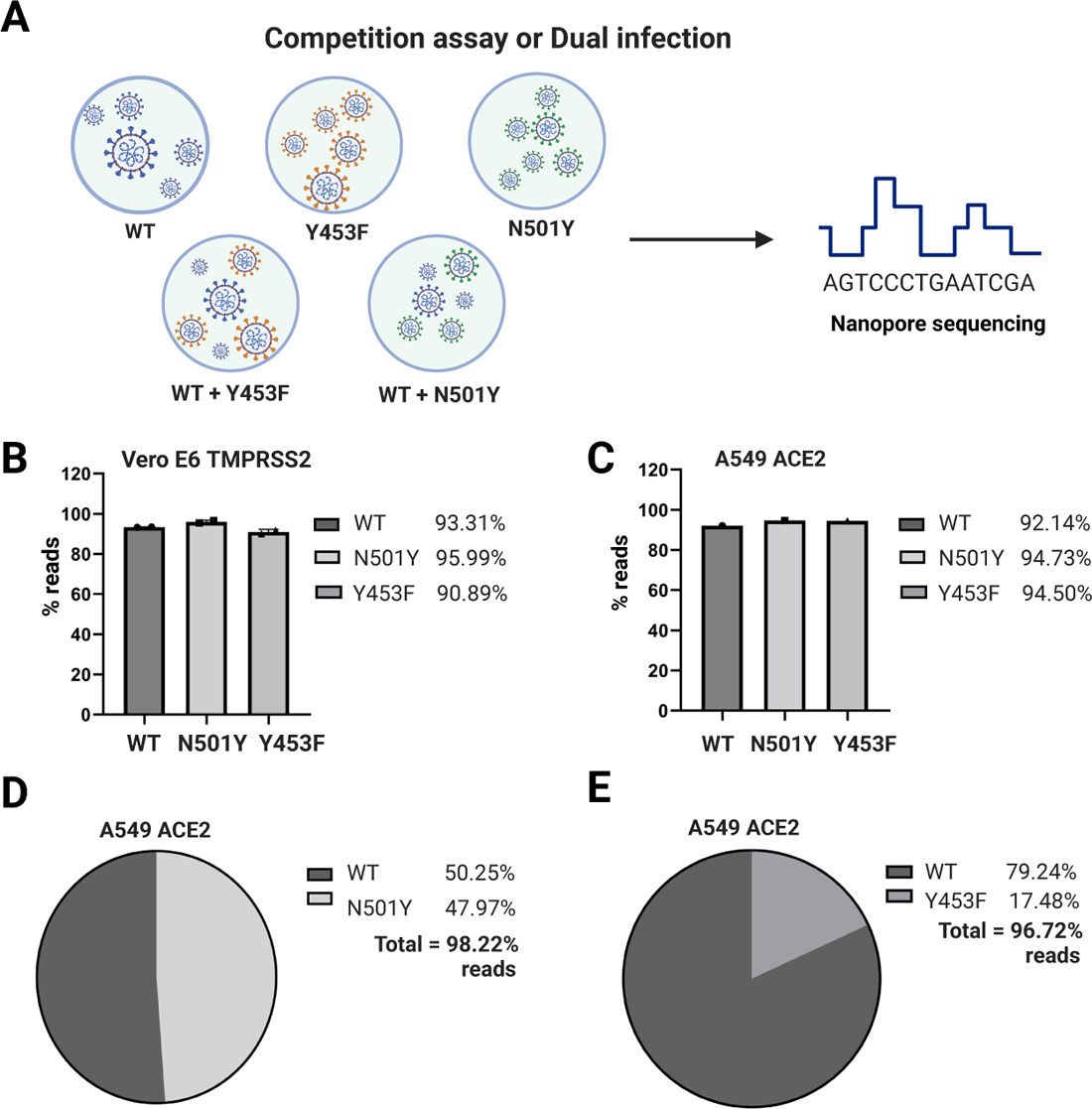
Competition assay for recombinant SARS-CoV-2 RBD spike mutants. **(A)** Experimental design for competition assay. **(B** and **C)** Controls for competition assay. **(B)** Vero E6 TMPRSS2 and **(C)** A549 ACE2 were infected at MOI 0.1 with rSARS-CoV-2 WT, rSARS-CoV-2 N501Y and rSARS-CoV-2 Y453F separately. After 48 hpi samples were collected and sequenced by Oxford Nanopore. **(D** and **E)** Competition assay for SARS-CoV-2 spike mutants. A549 were infected with rSARS-CoV-2 WT plus rSARS-CoV-2 N501Y and rSARS-CoV-2 WT plus rSARS-CoV-2 Y453F at a MOI 0.1. After 48 hpi samples coming from cells infected with **(D)** WT+N501Y or **(E)** WT+Y453F were collected and sequenced by Oxford Nanopore. Percentage of reads corresponding to the respective rSARS-CoV-2 mutant sequence. Vero E6 TMPRSS2 *n* = 2 independent experiment with two duplicates. A549 ACE2 *n* = 1 independent experiment with two duplicates.

**Supplementary Figure 8.**
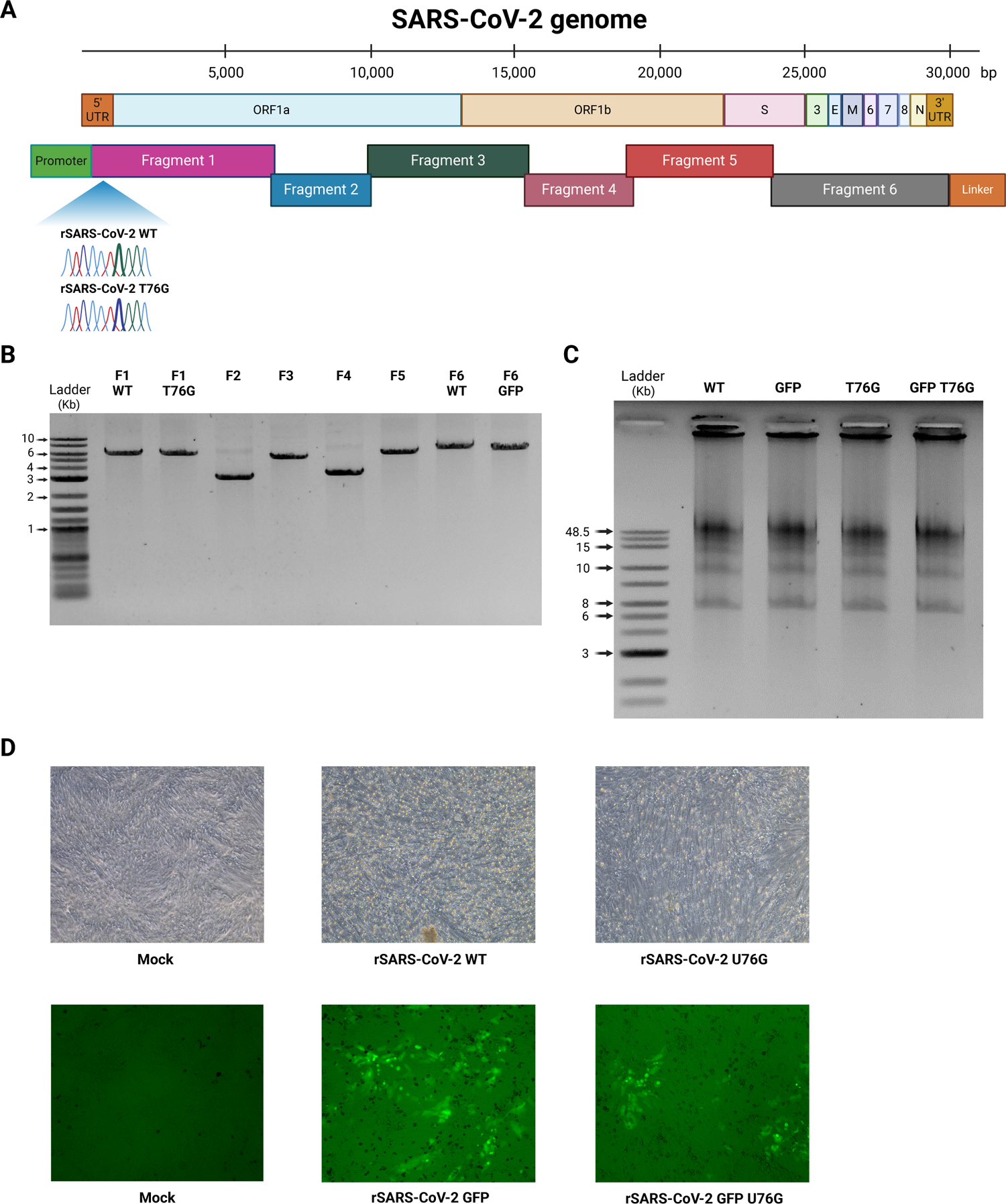
Design and generation of SARS-CoV-2 5’UTR mutant. **(A)** Schematic representation of SARS-CoV-2 genome and mutation T76G. **(B)** DNA fragments of SARS-CoV-2. PCR products of 8 fragments of SARS-CoV-2, including fragment 1 with T76G mutation. **(C)** Assembly of complete DNA SARS-CoV-2. An equimolar mixture of 6 DNA fragments is assembled in the 30 kb SARS-CoV-2 genome, in order to generate different recombinant SARS-CoV-2. **(D)** Infection using recombinant SARS-CoV-2 U76G mutants. After DNA transfection, supernatants were collected to infect a monolayer of Vero E6 TMPRSS2 cells, after 2 days CPE was showed for rSARS-CoV-2 WT and rSARS-CoV-2 U76G; or reporter GFP was expressed in the case of rSARS-CoV-2 GFP and rSARS-CoV-2 GFP U76G.

## Bibliography

1. Hu, B., Guo, H., Zhou, P. & Shi, Z. L. Characteristics of SARS-CoV-2 and COVID-19. Nat. Rev. Microbiol. 19, 141–154 (2021).

2. Brant, A. C., Tian, W., Majerciak, V., Yang, W. & Zheng, Z. M. SARS-CoV-2: from its discovery to genome structure, transcription, and replication. Cell and Bioscience vol. 11 1–17 (2021).

3. Lan, T. C. T. et al. Secondary structural ensembles of the SARS-CoV-2 RNA genome in infected cells. Nat. Commun. 13, 1–14 (2022).

4. Wang, C., Horby, P. W., Hayden, F. G. & Gao, G. F. A novel coronavirus outbreak of global health concern. Lancet 395, 470–473 (2020).

5. Wang, D. et al. The SARS-CoV-2 subgenome landscape and its novel regulatory features. Mol. Cell 81, 2135–2147.e5 (2021).

6. Duarte, C. M. et al. Rapid evolution of SARS-CoV-2 challenges human defenses. Sci. Rep. 12, 1–8 (2022).

7. Harvey, W. T. et al. SARS-CoV-2 variants, spike mutations and immune escape. Nat. Rev. Microbiol. 19, (2021).

8. Singh, J., Pandit, P., McArthur, A. G., Banerjee, A. & Mossman, K. Evolutionary trajectory of SARS-CoV-2 and emerging variants. Virol. J. 18, 1–21 (2021).

9. Thakur, S. et al. SARS-CoV-2 Mutations and Their Impact on Diagnostics, Therapeutics and Vaccines. Front. Med. 9, (2022).

10. Sonnleitner, S. T. et al. Cumulative SARS-CoV-2 mutations and corresponding changes in immunity in an immunocompromised patient indicate viral evolution within the host. Nat. Commun. 13, (2022).

11. Shannon, A. et al. Rapid incorporation of Favipiravir by the fast and permissive viral RNA polymerase complex results in SARS-CoV-2 lethal mutagenesis. Nat. Commun. 11, 1–9 (2020).

12. Polack, F. P. et al. Safety and Efficacy of the BNT162b2 mRNA Covid-19 Vaccine. N. Engl. J. Med. 383, 2603–2615 (2020).

13. Baden, L. R. et al. Efficacy and Safety of the mRNA-1273 SARS-CoV-2 Vaccine. N. Engl. J. Med. 384, 403–416 (2021).

14. Voysey, M. et al. Single-dose administration and the influence of the timing of the booster dose on immunogenicity and efficacy of ChAdOx1 nCoV-19 (AZD1222) vaccine: a pooled analysis of four randomised trials. *Lancet (London*, England) 397, 881–891 (2021).

15. Stobart, C. C. & Moore, M. L. RNA virus reverse genetics and vaccine design. Viruses vol. 6 2531–2550 (2014).

16. Almazán, F. et al. Coronavirus reverse genetic systems: Infectious clones and replicons. Virus Res. (2020).

17. Wang, W., Peng, X., Jin, Y., Pan, J. A. & Guo, D. Reverse genetics systems for SARS-CoV-2. J. Med. Virol. 94, 3017–3031 (2022).

18. Thi Nhu Thao, T., et al. Rapid reconstruction of SARS-CoV-2 using a synthetic genomics platform. Nature 582, 561–565 (2020).

19. Ricardo-Lax, I. et al. Replication and single-cycle delivery of SARS-CoV-2 replicons. Science (80-.). 374, 1099–1106 (2021).

20. Fahnøe, U. et al. Versatile SARS-CoV-2 Reverse-Genetics Systems for the Study of Antiviral Resistance and Replication. Viruses 14, (2022).

21. Beitzel, B., Hulseberg, C. E. & Palacios, G. Reverse genetics systems as tools to overcome the genetic diversity of Lassa virus. Curr. Opin. Virol. 37, 91–96 (2019).

22. Hou, Y. J. et al. SARS-CoV-2 Reverse Genetics Reveals a Variable Infection Gradient in the Respiratory Tract. Cell 182, 429–446.e14 (2020).

23. Ye, C. et al. Rescue of SARS-CoV-2 from a single bacterial artificial chromosome. MBio 11, 1–10 (2020).

24. Edmonds, J. et al. A Novel Bacterium-Free Method for Generation of Flavivirus Infectious DNA by Circular Polymerase Extension Reaction Allows Accurate Recapitulation of Viral Heterogeneity. J. Virol. 87, 2367–2372 (2013).

25. Neumann, G., Fujii, K., Kino, Y. & Kawaoka, Y. An improved reverse genetics system for influenza A virus generation and its implications for vaccine production. Proc. Natl. Acad. Sci. U. S. A. 102, 16825–9 (2005).

26. Nogales, A. & Martínez-Sobrido, L. Reverse genetics approaches for the development of influenza vaccines. Int. J. Mol. Sci. 18, 1–26 (2017).

27. Jung, E. J., Lee, K. H. & Seong, B. L. Reverse genetic platform for inactivated and live-attenuated influenza vaccine. Exp. Mol. Med. 42, 116–121 (2010).

28. Taniguchi, T., Palmieri, M. & Weissmann, C. Qβ DNA-containing hybrid plasmids giving rise to Qβ phage formation in the bacterial host. Nature 274, 223–228 (1978).

29. Racaniello, V. R. & Baltimore, D. Cloned Poliovirus Complementary DNA Is Infectious in Mammalian Cells. Science (80-.). 214, 912–912 (1981).

30. Pleschka, S. et al. A plasmid-based reverse genetics system for influenza A virus. J. Virol. 70, 4188–4192 (1996).

31. Almazán, F. et al. Engineering the largest RNA virus genome as an infectious bacterial artificial chromosome. Proc. Natl. Acad. Sci. U. S. A. 97, 5516–5521 (2000).

32. Wakita, T. et al. Production of infectious hepatitis C virus in tissue culture from a cloned viral genome. Nat. Med. 11, 791–796 (2005).

33. Kanai, Y. et al. Entirely plasmid-based reverse genetics system for rotaviruses. Proc. Natl. Acad. Sci. U. S. A. 114, 2343–2348 (2017).

34. Tischer, B. K. & Kaufer, B. B. Viral bacterial artificial chromosomes: Generation, mutagenesis, and removal of mini-F sequences. J. Biomed. Biotechnol. 2012, (2012).

35. Kouprina, N. et al. Segments missing from the draft human genome sequence can be isolated by transformation-associated recombination cloning in yeast. EMBO Rep. 4, 257–262 (2003).

36. Marschall, P., Malik, N. & Larin, Z. Transfer of YACs up to 2.3 Mb intact into human cells with polyethylenimine. Gene Ther. 6, 1634–1637 (1999).

37. Rihn, S. J. et al. A plasmid DNA-launched SARS-CoV-2 reverse genetics system and coronavirus toolkit for COVID-19 research. PLoS Biol. 19, e3001091 (2021).

38. Xie, X. et al. Engineering SARS-CoV-2 using a reverse genetic system. Nat. Protoc. (2021) doi:10.1038/s41596-021-00491-8.

39. Ju, X. et al. A novel cell culture system modeling the SARS-CoV-2 life cycle. PLoS Pathog. 17, 1–23 (2021).

40. Shi, P.-Y. et al. An Infectious cDNA Clone of SARS-CoV-2. Cell Host Microbe 27, 841– 848 (2020).

41. Torii, S. et al. Establishment of a reverse genetics system for SARS-CoV-2 using circular polymerase extension reaction. Cell Rep. 35, (2021).

42. Amarilla, A. A. et al. A versatile reverse genetics platform for SARS-CoV-2 and other positive-strand RNA viruses. Nat. Commun. 12, (2021).

43. Mélade, J. et al. A simple reverse genetics method to generate recombinant coronaviruses. EMBO Rep. 23, 1–14 (2022).

44. Setoh, Y. X., et al. De Novo Generation and Characterization of New Zika Virus Isolate Using Sequence Data from a Microcephaly Case. mSphere 2, (2017).

45. Sambrook, J. & Russell, D. W. Molecular Cloning: A Laboratory Manual. Third Edition. United States: Cold Spring Harbor Laboratory Press. (2001).

46. Marillonnet, S. & Grützner, R. Synthetic DNA Assembly Using Golden Gate Cloning and the Hierarchical Modular Cloning Pipeline. Curr. Protoc. Mol. Biol. 130, 1–33 (2020).

47. Setoh, Y. X. et al. Systematic analysis of viral genes responsible for differential virulence between American and Australian West Nile virus strains. J. Gen. Virol. 96, 1297–1308 (2015).

48. Carabelli, A. M. et al. SARS-CoV-2 variant biology: immune escape, transmission and fitness. Nat. Rev. Microbiol. 21, 162–177 (2023).

49. Markov, P. V et al. The evolution of SARS-CoV-2. Nat. Rev. Microbiol. 21, 361–379 (2023).

50. Bayarri-Olmos, R. et al. The SARS-CoV-2 Y453F mink variant displays a pronounced increase in ACE-2 affinity but does not challenge antibody neutralization. J. Biol. Chem. 296, 100536 (2021).

51. Motozono, C. et al. SARS-CoV-2 spike L452R variant evades cellular immunity and increases infectivity. Cell Host Microbe 29, 1124–1136.e11 (2021).

52. Liu, Y. et al. The N501Y spike substitution enhances SARS-CoV-2 infection and transmission. Nature 602, 294–299 (2022).

53. Starr, T. N. et al. Deep Mutational Scanning of SARS-CoV-2 Receptor Binding Domain Reveals Constraints on Folding and ACE2 Binding. Cell 182, 1295–1310.e20 (2020).

54. Zahradník, J. et al. SARS-CoV-2 variant prediction and antiviral drug design are enabled by RBD in vitro evolution. Nat. Microbiol. 6, 1188–1198 (2021).

55. Tian, F. et al. N501y mutation of spike protein in sars-cov-2 strengthens its binding to receptor ace2. Elife 10, 1–17 (2021).

56. Han, P. et al. Molecular insights into receptor binding of recent emerging SARS-CoV-2 variants. Nat. Commun. 12, (2021).

57. Barton, M. I. et al. Effects of common mutations in the sars-cov-2 spike rbd and its ligand the human ace2 receptor on binding affinity and kinetics. Elife 10, 1–19 (2021).

58. Cui, J., Li, F. & Shi, Z. L. Origin and evolution of pathogenic coronaviruses. Nat. Rev. Microbiol. 17, 181–192 (2019).

59. Hoffmann, M. et al. SARS-CoV-2 Cell Entry Depends on ACE2 and TMPRSS2 and Is Blocked by a Clinically Proven Protease Inhibitor. Cell 181, 271–280.e8 (2020).

60. Koch, J. et al. TMPRSS2 expression dictates the entry route used by SARS-CoV-2 to infect host cells. EMBO J. 40, 1–20 (2021).

61. Steuten, K. et al. Challenges for Targeting SARS-CoV-2 Proteases as a Therapeutic Strategy for COVID-19. ACS Infect. Dis. 7, 1457–1468 (2021).

62. Chang, C. W. et al. A Newly Engineered A549 Cell Line Expressing ACE2 and TMPRSS2 Is Highly Permissive to SARS-CoV-2, Including the Delta and Omicron Variants. Viruses 14, (2022).

63. Kim, Y. et al. Trypsin enhances SARS-CoV-2 infection by facilitating viral entry. Arch. Virol. 167, 441–458 (2022).

64. Haywood, A. M. Virus receptors: binding, adhesion strengthening, and changes in viral structure. J. Virol. 68, 1–5 (1994).

65. Kim, D. et al. The Architecture of SARS-CoV-2 Transcriptome ll ll Resource The Architecture of SARS-CoV-2 Transcriptome. Cell 181, 914–921.e10 (2020).

66. Kwon, K. & Beckett, D. Function of a conserved sequence motif in biotin holoenzyme synthetases. Protein Sci. 9, 1530–1539 (2000).

67. Wang, B., Svetlov, D. & Artsimovitch, I. NMPylation and de-NMPylation of SARS-CoV-2 nsp9 by the NiRAN domain. Nucleic Acids Res. 49, 8822–8835 (2021).

68. Longhi, S. et al. The severe acute respiratory syndrome-coronavirus replicative protein nsp9 is a single-stranded RNA-binding subunit unique in the RNA virus world. Proc. Natl. Acad. Sci. (2004) doi:10.1073/pnas.0307877101.

69. El-kamand, S. et al. A distinct ssDNA / RNA binding interface in the Nsp9 protein from SARS-CoV-2. Proteins 90, (2022).

70. Schmidt, N. et al. SND1 binds SARS-CoV-2 negative-sense RNA and promotes viral RNA synthesis through NSP9 ll ll SND1 binds SARS-CoV-2 negative-sense RNA and promotes viral RNA synthesis through NSP9. Cell 4834–4850 (2023) doi:10.1016/j.cell.2023.09.002.

71. Park, G. J. et al. The mechanism of RNA capping by SARS-CoV-2. Nature 609, 793– 800 (2022).

72. Small, G. I. et al. Structural and functional insights into the enzymatic plasticity of the SARS-CoV-2 NiRAN domain. Mol. Cell 83, 3921–3930.e7 (2023).

73. Furusawa, Y., Yamayoshi, S. & Kawaoka, Y. The accuracy of reverse genetics systems for SARS-CoV-2: Circular polymerase extension reaction versus bacterial artificial chromosome. *Influenza Other Respi*. Viruses 17, 2–5 (2023).

74. Ren, W. et al. Mutation Y453F in the spike protein of SARSCoV-2 enhances interaction with the mink ACE2 receptor for host adaption. PLoS Pathog. 17, 1–21 (2021).

75. Zhou, J. et al. Mutations that adapt SARS-CoV-2 to mink or ferret do not increase fitness in the human airway. Cell Rep. 38, (2022).

76. Lu, L. et al. The impact of spike N501Y mutation on neutralizing activity and RBD binding of SARS-CoV-2 convalescent serum. EBioMedicine 71, (2021).

77. Collier, D. A. et al. Sensitivity of SARS-CoV-2 B.1.1.7 to mRNA vaccine-elicited antibodies. Nature 593, 136–141 (2021).

78. Matsuyama, S. et al. Enhanced isolation of SARS-CoV-2 by TMPRSS2-expressing cells. Proc. Natl. Acad. Sci. U. S. A. 117, 7001–7003 (2020).

79. Tang, H. et al. Characterization of SARS-CoV-2 Variants N501Y.V1 and N501Y.V2 Spike on Viral Infectivity. Front. Cell. Infect. Microbiol. 11, 1–10 (2021).

80. Bloom, J. D. & Neher, R. A. Fitness effects of mutations to SARS-CoV-2 proteins. Virus Evol. 9, (2023).

81. Starr, T. N. et al. Shifting mutational constraints in the SARS-CoV-2 receptor-binding domain during viral evolution. Science (80-.). 377, 420–424 (2022).

82. Moulana, A. et al. Compensatory epistasis maintains ACE2 affinity in SARS-CoV-2 Omicron BA.1. Nat. Commun. 13, (2022).

83. Korber, B. et al. Tracking Changes in SARS-CoV-2 Spike: Evidence that D614G Increases Infectivity of the COVID-19 Virus. Cell 182, 812–827.e19 (2020).

84. Volz, E. et al. Assessing transmissibility of SARS-CoV-2 lineage B.1.1.7 in England. Nature 593, 266–269 (2021).

85. Yan, L. et al. Cryo-EM Structure of an Extended SARS-CoV-2 Replication and Transcription Complex Reveals an Intermediate State in Cap Synthesis. Cell 184, 184–193.e10 (2021).

86. Yan, L. et al. A mechanism for SARS-CoV-2 RNA capping and its inhibition by nucleotide analog inhibitors. Cell 185, 4347–4360.e17 (2022).

